# Flexible reorientation of conserved neural dynamics underlies grasp control

**DOI:** 10.64898/2026.09.04.749537

**Authors:** Zulfar Ghulam-Jelani, Matthew T. Kaufman

**Affiliations:** Committee on Computational Neuroscience; Department of Organismal Biology and Anatomy; Neuroscience Institute, The University of Chicago, Chicago, IL 60637 USA; NSF-Simons National Institute for Theory and Mathematics in Biology

## Abstract

Grasping objects is a complex behavior requiring high-dimensional hand control and dynamic interaction with the environment, yet its neural mechanisms remain poorly understood. Dynamical systems approaches have provided key insights into how neural populations generate control signals for arm movements such as reaching and cycling but have proven insufficient to account for neural population activity during grasping. Here, we re-analyzed multi-area neural recordings from rhesus monkeys performing a reach-to-grasp task to many objects and found two key population-level features. First, the component of the population trajectory that was common to all grasps formed a small angle with the subspace containing grasp condition-specific (object) tuning. Second, the neural activity was well fitted by a recent, flexible dynamical model developed for reaching, called location-dependent rotations (LDR). Neural activity in M1 and F5 during grasping exhibited conserved population-level rotational frequencies, but the state-space planes in which these rotations occurred varied systematically with grasp condition. The rotational center (the “location”) and the orientation of the rotations related both to each other and to the kinematics of movement. While these rotations were reoriented in high dimensional space, reflecting the high-dimensional nature of hand control, the extent of their tilt into additional dimensions were small, in contrast to reaching. This structure may reflect the nature of grasping as modulations of an overall open-close motif. Together, these results indicate that the LDR dynamics framework applies to grasping as well as reaching and provides an entry point to understanding how grasp commands are generated.

## INTRODUCTION

Grasping objects with the hand is a fundamental component of primate behavior. Despite its apparent ease in everyday life, grasping requires the coordinated control of many joints and muscles while integrating visual, proprioceptive, and tactile information in real time. To generate appropriate motor output, the nervous system must transform high-dimensional sensory information about object identity, shape, and location into precisely timed patterns of muscle activity that account for the complex biomechanics of the hand. Although substantial progress has been made in understanding the neural control of reaching movements^1–10^ and aspects of hand representation^11–13^, the neural computations underlying grasp remain poorly understood.

As a critical region in the nervous system for voluntary motor control, the motor cortex provides a window into the neural computations that occur at the final stages of cortical processing before movement execution. Classical views of motor cortex function proposed that individual neurons directly encode specific movement parameters, such as direction^14,15^, speed^3,16^, force^17–19^, amplitude^20,21^, or posture^22,23,13,24^. However, motor cortex neurons exhibit rich, time-varying activity patterns that are not fully captured by representational models that directly relate neural activity to kinematics^25–27^. Instead, a growing body of work has shown that a major component of motor cortex population activity can be described using the dynamical systems framework^27,5–7,28,8,29,10^, which characterizes how neural activity evolves over time according to mathematical rules. From this perspective, the motor cortex acts as a transiently autonomous dynamical system, perhaps in combination with consistent inputs, in which population activity generates subsequent activity patterns consistent with a role as a pattern generator^25,29–33^.

Most dynamical analyses of the motor cortex have primarily focused on low-dimensional behaviors, such as reaching, and have relied on simplifying assumptions to maintain mathematical tractability^5^. In these studies, neural dynamics were frequently characterized as “rotational”, with population activity tracing smooth, planar trajectories that orbit in state space. Such models assume a limited set of characteristic frequencies and fixed rotational planes, implying that the same neural subspaces are engaged across movements and that a small number of activity patterns dominate motor execution. These approaches successfully captured important aspects of reaching dynamics^5^. While these assumptions have proven useful, they accounted for only a fraction of movement-specific variance^5^, describe neural activity well only in restricted domains^34^, and notably fail to capture neural population activity during grasping^35^.

Recent work has challenged these assumptions by introducing a more flexible framework for characterizing motor cortical dynamics^10^. In the context of reaching, they showed that the subspace containing neural dynamics varies substantially and systematically across movements. In this Location-Dependent Rotations (LDR) model, dynamics are still rotational, with all reaches sharing a small number of fixed frequencies, but the embedding of these rotations in neural space varies by movement – allowing neurons to express different amplitudes and phases of each frequency for different reaches. This approach captures nearly all motor cortex activity during reaching and found tight relationships between reach kinematics and single-trial spiking. Importantly, it suggests that a representational-like signal is encoded in the overall “location” of neural activity within state space, which orients dynamics that then produce the time-varying motor commands.

Grasping has presented an even greater challenge both for controlling the hand and for analysis. Compared to reaching, grasp control involves many more joints and degrees of freedom, complex object-hand interactions, and extensive sensory feedback integration. Consistent with this complexity, prior work has shown that grasp-related neural population activity differs markedly from activity observed during reaching^35^. During grasp, the condition-independent component of activity is not orthogonal to the tuned activity^36^; and neural trajectories appear substantially more ‘tangled’^28^ than those associated with reaching, suggesting that grasp-related activity may not be well-described by a single low-dimensional autonomous flow field. Moreover, the limited improvements in grasp decoding performance achieved with sophisticated dynamical latent-variable approaches such as LFADS^37^ suggest that this complexity cannot be attributed solely to neural noise. Together, these observations indicate that the assumptions underlying existing dynamical models of reaching may not generalize to grasp.

Here, we applied the LDR framework and other dynamical systems analysis tools to neural population activity recorded from the grasp network including primary motor cortex (M1), grasp-related premotor cortex (F5), and the anterior intraparietal area (AIP) during a reach-to-grasp behavior. These areas are thought to collectively transform object-related visual information (in AIP) into motor commands for the hand (in F5 and M1)^38–41^. We show that grasp obeys LDR-like dynamics, in a somewhat different quantitative regime. Unlike in reaching, we replicated the finding that a strong component common to all conditions was present in the same dimensions as the tuned components, helping to mask the structure in those tuned components^36^. Second, using the extensive kinematic quantification in these data, we found that a large fraction of the neural activity correlated tightly with kinematics, suggesting that much of the complexity may be due to how many distinct joints must be controlled. Third, LDR described a sizable fraction of the data: oscillatory frequencies were conserved across grasps but their embedding in neural space varied systematically with the grasp kinematics. This suggests a shared mechanism with reaching in the dynamics of M1 especially. Finally, we found that the variation in state space location was smaller for grasp than reach, and the accompanying reorientation in rotational planes remained high-dimensional but varied through smaller angles. This difference is consistent with grasp-related dynamics being structured to enable control of many joints, with high-dimensional but smaller modulation of an otherwise shared open-close component.

## RESULTS

In this work, we re-analyzed data from previously published experiments^38–41^ in which two rhesus monkeys (M and Z) performed a delayed reach-to-grasp task. The monkeys were presented with objects of various shapes and sizes always at the same location, evoking 44 distinct grasp types (Fig. 1a), which we refer to as “conditions”. After briefly viewing an object, the lights were turned off, and following a Go cue, they reached, grasped, and briefly lifted the object. Only successful trials were analyzed, and activity from each cortical area and monkey was analyzed separately. Full arm and hand joint kinematics were recorded and aligned to object “contact”, which throughout this paper refers to the time when the hand crossed a spatial threshold just short of the object location, before actual contact (which was not instrumented). Single- and multi-unit neural data (analyzed together here) were recorded from M1 (61-160 units), F5 (73-110 units), and AIP (56-82 units). To ensure reliable behavioral measurements, trials with inconsistent joint angle trajectories were excluded based on trial-to-trial kinematic variability (Fig. 1b, orange; Methods). The remaining trials exhibited stereotyped, condition-specific joint trajectories, with the objects producing reliably different kinematic patterns across hand joints (Fig. 1b, other colors). Throughout this paper, condition traces are colored according to the object color scheme in Fig. 1a, with lighter colors indicating smaller objects and darker colors indicating larger objects.

**Figure 1:**
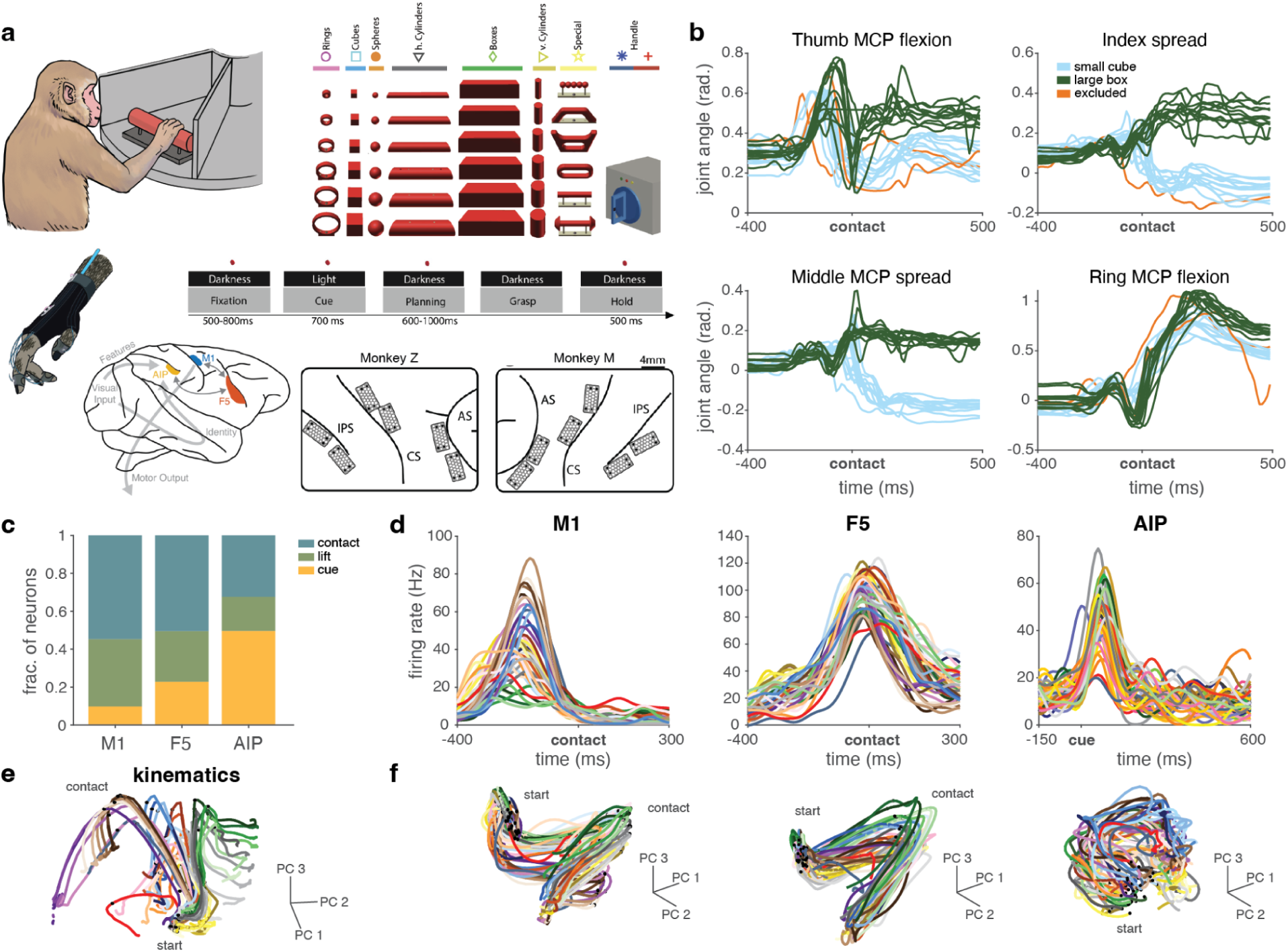
Reach-to-grasp task data. **a.** Schematic of the delayed reach-to-grasp task, from Schaffelhofer et al., 2015, Menz et al., 2015, and Michaels et al., 2020. **b.** Kinematics for representative joints and grasp conditions. Each trace shows one trial; excluded trials are shown in orange. **c.** Distribution of the temporal alignments that produced maximal peak firing rates across cortical areas, used to select the optimal alignment for each area. **d.** Example peri-event time histograms (PETHs) across the grasping network, aligned to the task events selected in **c** and colored by grasp condition using the same color scheme as in **a**. **e.** PCA trajectories of joint angle kinematics in low-dimensional state space, colored by grasp condition. Dots indicate start and contact. **f.** PCA trajectories of population activity in M1, F5, and AIP neural state space, colored by grasp condition. Dots indicate start and contact.

Neural activity was aligned to three different events: the onset of the brief viewing window (“cue”), the hand lifting from the hand rest, and object contact as defined above. For each unit and condition, we identified the event for which data alignment produced the maximal peak firing rate and report the fraction of unit-condition pairs assigned to each event. Consistent with prior results^38^, most units in M1 and F5 were maximally active when aligned to object contact, whereas AIP neurons were more strongly cue-modulated (Fig. 1c). Subsequent analyses are therefore shown aligned to object contact for M1 and F5 and to cue onset for AIP. Neural activity was smoothed with a Gaussian kernel (s.d. 30 ms) and trial-averaged within condition where applicable to estimate firing rates for subsequent analyses.

Examination of peri-event time histograms (PETHs) for an example unit in each area revealed that activity tended to have a similar structure over time across grasp conditions (Fig. 1d; more examples in Supp. Fig. 1). This shared temporal profile might reflect common aspects of the task across grasp conditions, such as reaching to a fixed spatial location and hand opening and closing movements occurring at roughly the same time, or a large common input (a “condition-independent signal”) that has been argued to trigger active dynamics and movement in reaching^42^. Although individual neurons showed condition-specific modulation, this tuning was typically smaller than the shared temporal structure of the responses. To avoid ascribing a particular function of the changes over time in the average activity of each neuron, and to acknowledge that the current shared temporal structure likely reflects a mixture of dynamics triggering, shared reach-related activity, and shared open-close grasp activity, we hereafter refer to it simply as the “common signal” for each neuron, or the “common trajectory” for the population.

### Population structure of grasp kinematics and neural activity

To first characterize the population structure of both hand kinematics and neural activity, we applied principal component analysis (PCA) to the trial-averaged data. PCA of joint angle time courses revealed well-separated trajectories corresponding to different grasp conditions (Fig. 1e), corresponding to reliably different hand configurations across objects^38,43,44^. In contrast, neural population activity in M1, F5, and AIP followed a largely shared, twisted-corkscrew manifold with modest separation between conditions that interacted with the corkscrew shape of the overall trajectory (Fig. 1f, second monkey in Supp. Fig. 1). These results confirm the presence of a large common signal in the neurons of each area, which interacts with the tuned structure.

To better understand whether and how these large common signals and the condition-specific (tuned) structure interact, we applied demixed principal component analysis (dPCA)^45^. In many previous tasks and cortical areas^45^, including M1 and dorsal premotor cortex during reaching^42^, dPCA has revealed the presence of a large condition-independent signal that is mostly orthogonal to the tuned structure. In the present grasp dataset, however, and consistent with other results in grasping^36^, dPCA did not reveal such orthogonalized structure. Despite successful variance demixing by dPCA (Fig. 2a,b), subspace analyses showed substantial geometric overlap between condition-independent (time) and condition-specific components. To quantify this geometry, we computed the principal angles between the condition-independent and condition-specific dPCA subspaces. Principal angles provide a direct measure of subspace geometry, where small angles indicate that there are dimensions of the two subspaces that form small angles with one another, whereas angles approaching 90° indicate that the subspaces are nearly orthogonal in their entirety. The principal angles between the condition-independent and condition-specific subspaces during grasping were modest, especially in M1 and F5 (M1: 13.4-18.2°, F5: 26.2-29.5°, AIP: 46.6-55.7°; range over monkeys). Although prior studies of reaching have described condition-independent and condition-specific activity as largely orthogonal^42^, this relationship has not, to our knowledge, been explicitly quantified geometrically. For comparison, we therefore performed the same analysis in M1 and PMd during reaching in the “maze” data that were previously used to study the condition-independent signal^42^, and found that the angles were substantially larger than in grasping (M1: 60.6-64.8°; PMd: 54.4-56.5°). Interestingly, though, even in the reaching data the principal angles were well less than orthogonal. Nevertheless, in contrast to reaching^42^ and cycling^28,46^ tasks, where condition-independent signals and condition-specific (tuned) activity occupy quite different subspaces, condition-independent signals in this grasping dataset resided in strongly overlapping subspaces with the condition-specific tuning. This overlap followed a gradient across areas, being weakest in AIP and progressively stronger in F5 and M1, indicating decreasing distinction between subspaces along the cortical hierarchy.

**Figure 2:**
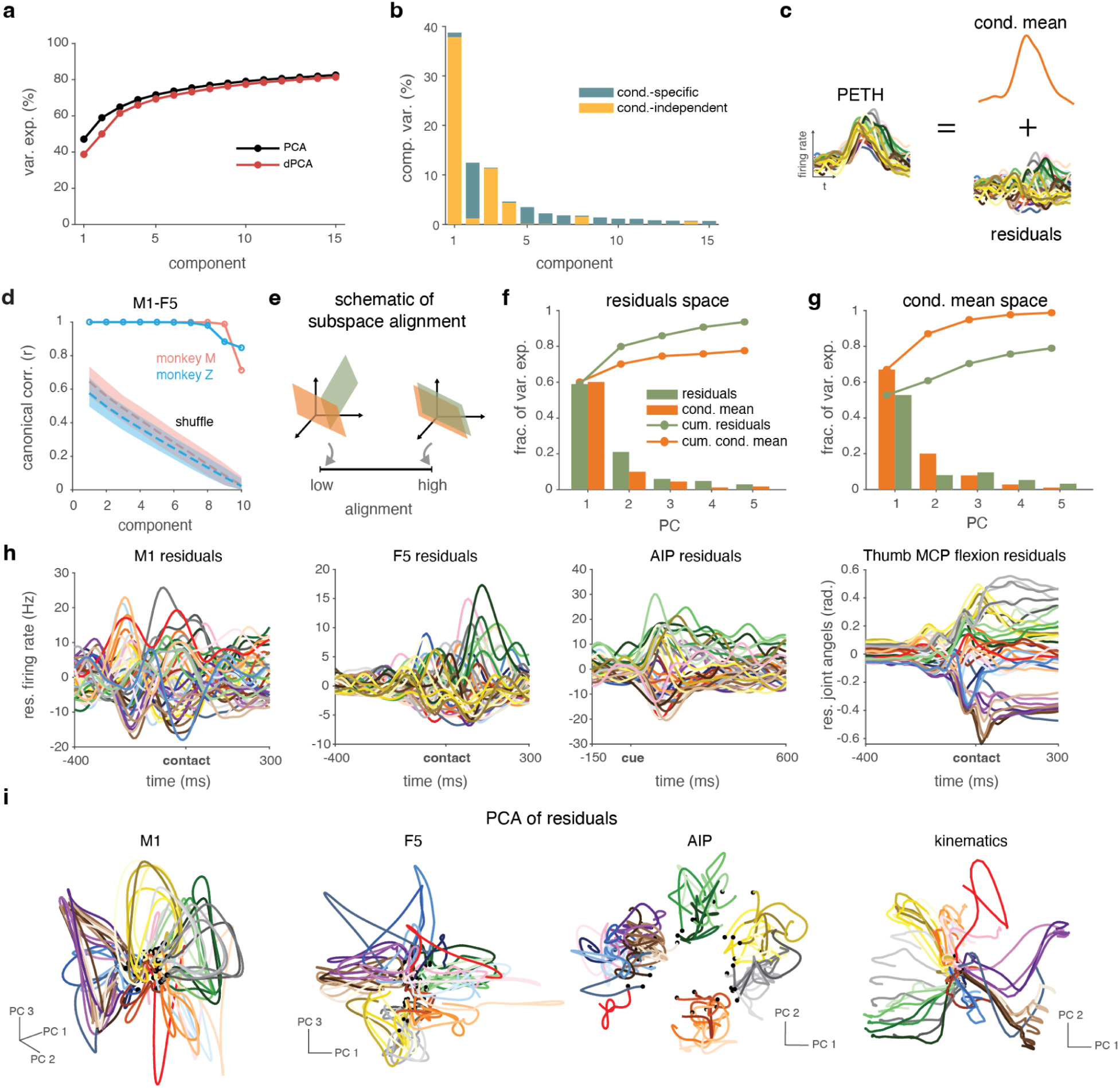
Overlap in condition-independent and grasp condition-specific activity. **a.** Cumulative variance explained by PCA (black) and dPCA (red) for M1 population activity. **b.** Variance explained by individual demixed principal components in M1. Bars indicate the proportion of total variance explained and are partitioned into condition-specific variance (blue) and condition-independent variance (yellow). Note that the basis is non-orthogonal and so the variance contained in the condition-independent and the condition-specific subspaces is partly shared; the total variance explained by a component may therefore be less than the sum of its marginalized parts. **c.** Schematic showing the condition-mean and residual components of a PETH. **d.** Shared condition-mean structure in M1 and F5 identified using CCA. Canonical correlations for the observed data are shown as solid lines, with permutation-based shuffle controls shown as dashed lines (mean; shading indicates 95% confidence interval). Colors denote individual monkeys. **e.** Schematic illustrating subspace alignment: trajectories are projected onto the PCs of a second set of trajectories; the captured variance in each PC is then normalized by the variance captured within the first set’s own corresponding PCs. **f.** Variance explained of the condition-mean (orange) and of the residual activity (green) in residuals space. **g.** Variance explained of the condition-mean and of the residual activity in condition-mean space. **h.** PETHs for example neurons (first three panels) or thumb metacarpal-phalangeal joint (last panel) after subtracting the condition-mean. Sinusoidal activity is apparent in the M1 and F5 examples. **i.** PCA trajectories of neural population activity for each area (M1, F5, AIP) and joint angle kinematics after subtraction of the condition-mean, colored by grasp condition.

### The common signal overlaps condition-specific activity

The geometric similarity between the condition-independent and condition-specific subspaces warranted further efforts to understand the relationship between the activity in each subspace. Continuing with dPCA, however, presented two problems: first, the basis produced by dPCA is non-orthogonal, introducing dependencies between the components; and second, the decoder axes found by dPCA can depend on choices of regularization and optimization objective. We therefore adopted a simpler and more direct approach to remove the common signal: we simply subtracted the across-condition mean (hereafter, “condition-mean”) for each time point for each unit (Fig. 2c). This procedure decomposes the activity into (i) a common trajectory reflecting structure shared across all grasps (whether due to reaching to a common target location, common open-close-related activity, dynamical initiation, or something else), and (ii) residuals / grasp-specific activity reflecting condition-specific deviations from the common trajectory. This residualization strategy is conceptually related to recent approaches that analyze residual neural activity after removing dominant shared signals^47^, although here the residuals reflect deviations across grasp conditions rather than trial-to-trial fluctuations used to estimate recurrent dynamics.

To assess whether the common trajectory’s structure was shared across cortical areas, we applied canonical correlation analysis (CCA) to the condition-mean trajectories from M1 and F5. We did not compare AIP because its activity was largely a function of the visual cue. The leading canonical dimensions showed almost perfect correlations between M1 and F5 (Fig. 2d), indicating that the condition-mean structure was highly similar across these two areas in the grasping network. For a baseline comparison, we repeated the analysis after randomly permuting the temporal order of the points in the F5 trajectories before smoothing. This permutation substantially reduced the canonical correlations, indicating that the observed alignment depended on the temporal structure of the trajectories rather than arising from chance or as an artifact of smoothing finite data.

To understand the relationship of the condition-mean component and condition-specific component within-area, we quantified the overlap between the condition-mean subspace and the condition-specific (residual) subspace using the alignment index, which measures how much variance from one subspace is captured by the principal components of the other^48,49^ (Fig. 2e). Assessed in either direction, variance was concentrated in the first few principal components and the same dominant dimensions captured variance in both, indicating that the highest-variance dimensions of the two subspaces were strongly overlapping (Fig. 2f,g, M1 shown; other areas and monkey in Supp. Fig. 2).

In neural state space, subtraction of the condition-mean revealed, as expected, grasp-specific trajectory structure in all three cortical areas that was obscured in the original PETHs (Fig. 2h; second monkey in Supp. Fig. 1d) and top PCs (Fig. 2i; second monkey in Supp. Fig. 1c). The condition-specific (residual) activity exhibited clear structure. In the PETHs, we observed clear difference in firing rates between objects, and also that firing rates of neurons in M1 and F5 exhibited strong sinusoidal patterns over time – similar to the activity patterns during reaching that first suggested the possibility of rotational dynamics (Fig. 2h; second monkey in Supp. Fig. 1d). For all three areas, the low-D trajectories for grasps to different sizes of the same object were grouped, and grasps to different objects were separated. For M1 and F5, grasps to different objects appeared to traverse largely different subspaces (Fig. 2i; second monkey in Supp. Fig. 1e). For AIP, the residuals had weak temporal structure and activity for different objects was simply stably segregated. Applying the same mean-subtraction procedure to the kinematic data similarly made the relationships between related movements clearer, while keeping different grasps well separated (Fig. 2h,i, kinematics; second monkey in Supp. Fig. 1d,e).

### Grasp exhibits dynamical structure before sensory feedback dominates cortical activity

Previous work, using a different task and recordings, argued that grasp-related activity is not consistent with M1 acting as a transiently autonomous dynamical system^35^. There, the neural data were poorly fit by a linear dynamical system, and population trajectories were tangled, indicating trajectories that are inconsistent with a smooth dynamical system in general. Here, using different recordings and a grasping task that included both many more objects as well as a reach to a shared location, we evaluated whether dynamical systems could fit the neural data in this context.

As a first test, we applied jPCA to test for the simplest version of planar rotational dynamics that were previously proposed for reaching^5^. This was performed with the typical procedure including condition-mean subtraction. Across M1, F5, and AIP, only 10-17% of the total variance was captured, and the resulting projections exhibited unconvincing rotations (Fig. 3a). This result argues that the simplest dynamical model, of rotational dynamics in a single plane for all grasp conditions, does not fit these data well.

**Figure 3:**
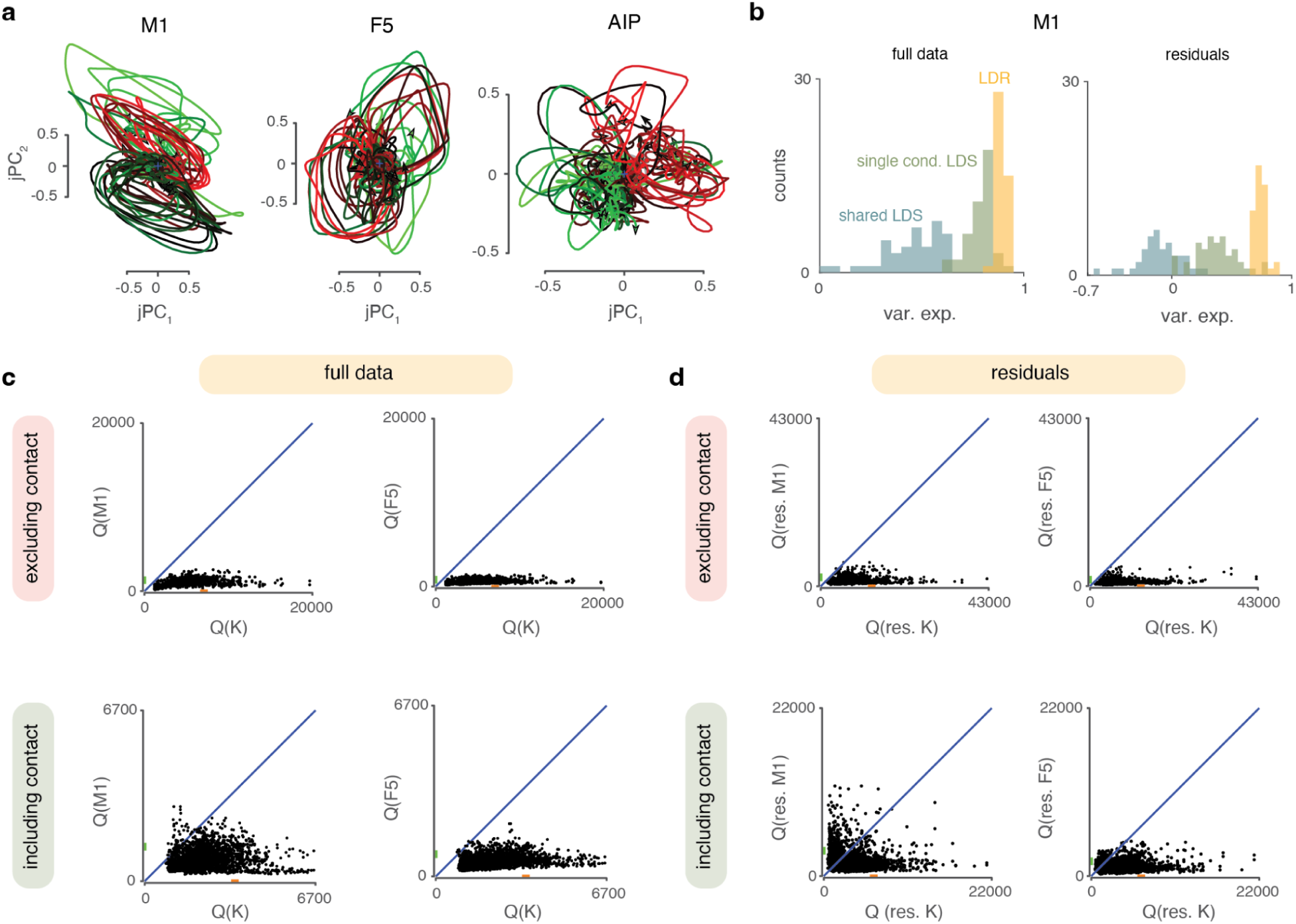
Testing dynamical systems properties in grasp-related neural activity. **a.** jPCA projections of trial-averaged neural population activity for individual grasp conditions in M1, F5, and AIP. **b.** Cross-validated population variance explained by a shared linear dynamical system (LDS) fit across conditions (blue), condition-specific LDSs fit separately to each grasp condition (green), and the location-dependent rotations (LDR) model (yellow). Left, fits to the full data; right, fits after subtracting the condition-mean. **c.** Trajectory tangling of full neural activity in M1 and F5 (Q(M1), Q(F5)) plotted against kinematic tangling (Q(K)) for analyses including only the pre-contact period (top) or including the full trial including object hold time (bottom). **d.** Same as **c** but computed using residualized neural activity (Q(res. M1), Q(res. F5)) and residualized kinematics (Q(res. K)) by subtracting the condition-mean.

We next asked whether the weak rotational structure reflected trajectories that were individually incompatible with linear dynamics (e.g., S-shaped trajectories). Alternatively, trajectories could be individually compatible with a linear dynamical system (LDS), but different conditions might obey different dynamics (e.g., rotational trajectories having different frequencies, or occupying entirely different planes). To do so, we fit LDSs to the data using either a single LDS shared across conditions (Fig. 3b, blue) or separate LDSs fit to each condition independently (Fig. 3b, green; F5 and second monkey in Supp. Fig. 3). For these analyses, we focused on M1 and F5 data. We performed these analyses both without condition-mean subtraction (Fig. 3b, left) and with it (Fig. 3b, right). The shared LDS explained a modest fraction of the variance of the common signal, but effectively none of the condition-specific activity (cross-validated fits with condition-mean subtraction were negative, at −9.8%; but 43.1% without condition-mean subtraction). This argues that the shared LDS fit explained only the common trajectory. The single-condition LDS fits performed appreciably better, explaining 78.8% of the total variance and 37.3% of the condition-specific variance (with cross-validation). This indicates that individual trajectories were at least partly compatible with linear dynamics, but that the planes and/or frequencies of rotations were not shared across conditions.

Linear dynamical systems impose strong constraints on neural trajectories. We therefore next asked whether grasp-related population activity could be described by a single set of smooth autonomous dynamics more generally, including potentially nonlinear dynamics. To do so, we computed trajectory tangling^28^, a measure of whether nearby points in neural state space evolve in similar directions, as required for a smooth dynamical system. High tangling occurs when trajectories nearly cross or diverge sharply, with nearby points evolving in different directions; whereas low tangling indicates that trajectories with different derivatives remain well separated in state space. Previous work has shown that during reaching and cycling behaviors, neural population activity in multiple cortical areas exhibits lower tangling than either muscle activity or movement kinematics^28^, consistent with the idea that cortical networks function as dynamical pattern generators while muscles are more strongly input-driven. In contrast, during grasping behaviors in which a robotic arm transported objects to the animal’s hand for grasping, neural activity in both motor and somatosensory cortices was as tangled as, or more tangled than, the corresponding hand kinematics^35^.

In the present dataset, which includes a reaching component prior to grasp that was similar for all conditions, we examined tangling with a variety of data preprocessing choices. Considering activity preceding contact, to ensure that the object-specific external feedback forces were excluded, we found that tangling was low in both M1 and F5 (Fig. 3c, top; second monkey in Supp. Fig. 3). Tangling was again low in M1 and F5 if we residualized the data by subtracting the condition-mean (Fig. 3d, top). This low tangling argues that the neural activity in these motor areas was consistent with a smooth dynamical system, whether or not we included the common trajectory. If we instead included activity from after object contact, tangling rose moderately, presumably due to the external sensory feedback inputs differing between conditions (Fig. 3c, bottom). Nevertheless, tangling remained modest for both M1 and F5. In the residualized data, however, if contact was included the level of tangling rose dramatically, especially in M1 (Fig. 3d, bottom). This suggests that the condition-specific aspects of the trajectories, which we would expect to mostly reflect the differing grasps, were highly impacted by object-specific feedback. Together, these results argue that in this task the grasp-related activity in M1 and F5 is consistent with transiently autonomous dynamics until object contact provides condition-specific external inputs.

### Temporal frequencies are conserved but the rotational planes vary by grasp condition

The firing rates of neurons in motor cortical areas during reaching exhibit multiphasic, quasi-sinusoidal responses^5^. Although multiphasic changes in firing rate were not immediately apparent in PETHs for grasp activity (Fig. 1d), subtracting the condition-mean made sinusoidal changes in neuron’s firing rates obvious (Fig. 2h; Supp. Fig. 1d). In initial work, the coordinated, sinusoidal changes in firing rates for reaching were described in terms of rotational dynamics: fixed frequencies of rotation occurring consistently within a conserved subspace^5^. More recent analyses have shown that these dynamics are better described by a more-flexible variant of this model, location-dependent rotations^10^. LDR preserves the axiom that rotational frequencies are consistent across movements but allows the plane (the subspace) of each rotation to vary by condition. Further, it relates these changes in rotational planes to condition-specific changes in the location of the rotational center’s location in state space, consistent with a globally curved ‘shape’ of the manifold orienting the rotations.

Given the combination of strong sinusoidal modulation of firing rate in grasp, the poor performance of a shared LDS in describing condition-specific activity here (Fig. 3b), the compatibility of individual grasp conditions with dynamical systems (Fig. 3c-d), the greater flexibility of the LDR model, and LDR’s better description of reaching data, we asked whether LDR might better describe grasp data than previous dynamical systems models. We therefore tested each aspect of LDR in turn. Analyses were performed on both the unmodified data and condition-mean subtracted (residualized) data wherever both were applicable.

We first asked whether rotational frequencies were conserved across grasp conditions. To do so, we fit low-D linear dynamical systems separately to each grasp condition (as in Fig. 3b, green; Methods) and examined the resulting eigenvalue spectra (Fig. 4a). Across all three cortical areas, we observed consistent eigenvalues corresponding to conserved rotational frequencies across conditions (Fig. 4b; second monkey in Supp. Fig. 4). In M1 and F5, dominant frequencies occurred at approximately 1 Hz and 2.5 Hz, whereas AIP exhibited a single dominant rotation near 1 Hz. To quantify this conservation, we computed the pairwise distances between same-mode eigenvalues in the complex plane and compared this distribution with the noise expected due to finite sampling in single conditions (Fig. 4c, M1 shown; other areas and monkey in Supp. Fig. 4). By this measure, the eigenvalues were as consistent as permitted by estimation noise, indicating preservation of rotational frequencies across grasp conditions.

**Figure 4:**
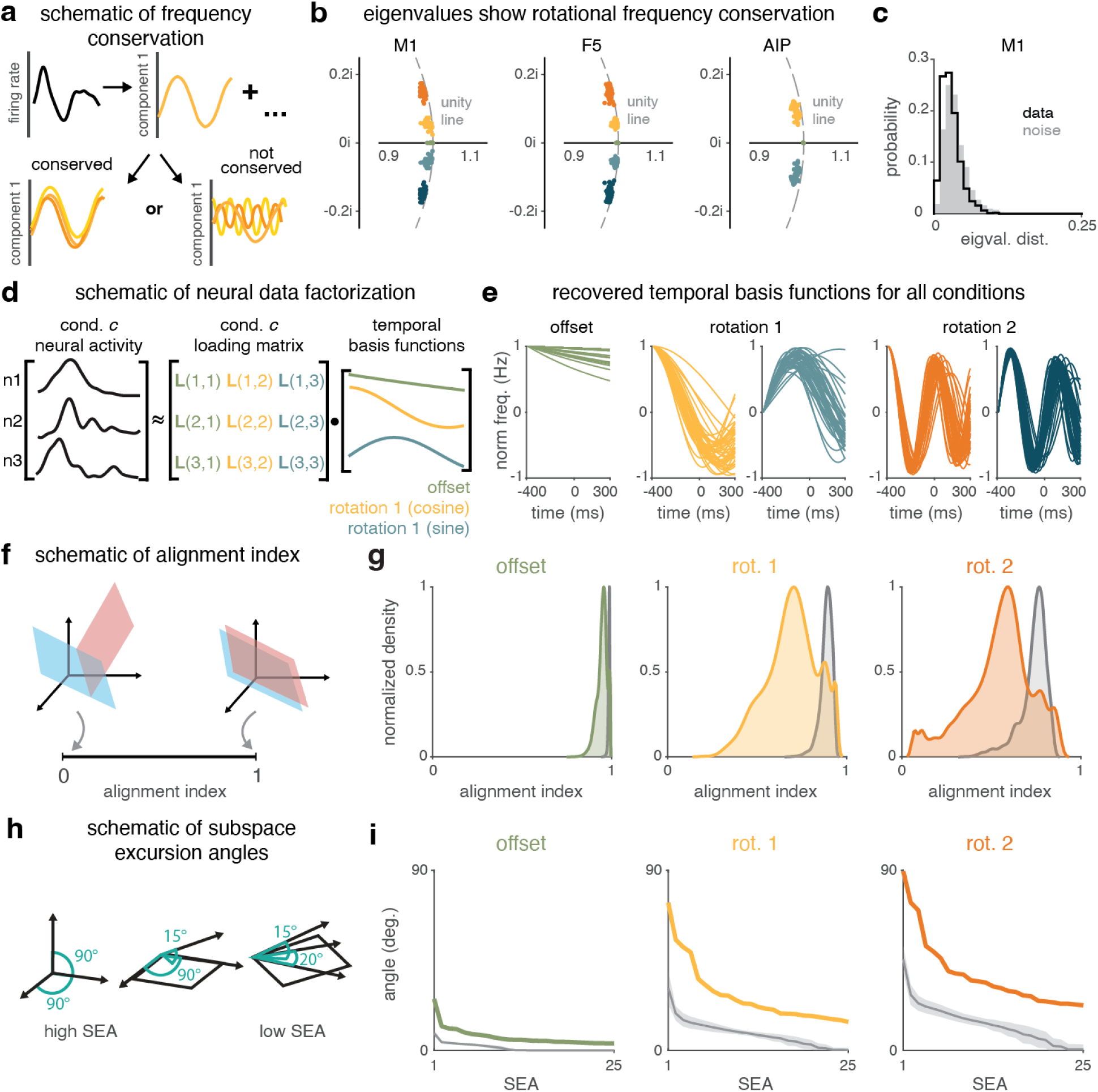
Conserved temporal frequencies are expressed through condition-specific geometry. **a.** Schematic illustrating the decomposition of firing rate activity to assess whether grasp conditions exhibit conserved temporal frequencies. **b.** Eigenvalues of the inferred dynamics for all grasp conditions. Clumped eigenvalues indicate that similar rotational frequencies were found across conditions. **c.** Eigenvalue distance between conditions for M1, computed as the mean distance between corresponding (closest-frequency) eigenvalues in the complex plane. Noise distribution computed by partitioning the set of trials into halves and computing eigenvalues from the two halves for the same condition separately. **d.** Schematic illustrating LDR neural data factorization. **e.** Recovered temporal basis functions across all conditions in M1 (data projected using loadings for a single component), aligned to contact. Version including object hold epoch shown. **f.** Schematic of the alignment index, which quantifies the overlap between pairs of rotational planes. **g.** Alignment indices between corresponding (same-frequency) rotational planes across pairs of grasp conditions (colored), compared with the distribution expected from estimation noise (gray) in M1. Distribution shown with kernel density estimation. **h.** Schematic illustrating subspace excursion angles (SEAs). **i.** SEAs between corresponding rotational planes across conditions (colored), along with angles expected from estimation noise (gray) in M1. For the noise distribution, the line indicates the mean; shaded regions indicate ±1 s.d.

This frequency conservation justifies applying the central decomposition of LDR, in which neural activity is factorized into two parts (Fig. 4d): (1) a set of temporal basis functions that encode conserved oscillatory frequencies shared across conditions (the rotations), and (2) condition-specific loadings that determine the magnitude and phase of each frequency in neural state space. As in reaching^10^, this factorization was performed by constructing a trial-averaged neuron * condition x time data matrix and applying singular value decomposition.

To assess how well this model fit the data, we first examined how cleanly isolated the different frequencies were from one another in state space. To do so, we asked how consistently the projected data could recover the temporal basis functions (Fig. 4e, M1 shown). In almost every case, the data produced well conserved frequencies, as expected. To quantify the fits, we then compared LDR’s cross-validated fit performance with the previous LDS fits (Fig. 3b, yellow). The LDR fits were substantially better than the LDS: 88.8% of M1’s total variance fitted without condition-mean subtraction (p<0.001 relative to single-condition LDS, Wilcoxon signed-rank test), and 74.1% with condition-mean subtraction (p<0.001 relative to single-condition LDS, Wilcoxon signed-rank test). Similar results were observed in F5 (Supp. Fig. 3).

We next asked how strongly the rotational planes varied in neural state space. To quantify subspace similarity, we computed the alignment index^48–50^, which measures how well the low-dimensional subspace traversed by one trajectory captures the variance in another trajectory. Values range from 0 (orthogonal subspaces) to 1 (identical subspaces; Fig. 4f). Same-frequency rotational planes showed significantly lower alignment across conditions than expected simply due to estimation noise (Fig. 4g, M1 shown; other areas and monkey in Supp. Fig. 4), indicating that the rotational planes indeed varied by condition. Nonetheless, alignments remained far from zero, indicating that the rotational planes were not entirely orthogonal across conditions.

To geometrically characterize the variation in rotational plane orientation, we computed Subspace Excursion Angles (SEAs)^10^, which quantify the largest series of angular steps that can be taken between subspaces while avoiding previously traversed subspaces (schematized in Fig. 4h). Intuitively, this measure asks not just how many dimensions the rotational planes vary in, but how tilted into each dimension different conditions were. Across all three cortical areas, SEAs for corresponding rotational planes were large and substantially exceeded values expected from estimation noise (number of angles >45°: M1, 4-6; F5, 2-7; AIP: 0-2; Fig. 4i, M1 shown; other areas and monkey in Supp. Fig. 4). When data were residualized to remove the common signal, the subspace alignments across conditions dropped further still, and the SEAs rose (Supp. Fig. 5). This finding is consistent with the grasp-related dynamics strongly resembling what was previously seen for reaching, but with a strong common signal that makes the conditions more similar. Note, however, that the residualized signals were smaller, so the expected alignment due to noise was also lower. In summary, together these results further support the finding of substantial variation in the orientation of the rotational planes through several dimensions.

### Reorientation of dynamical subspaces is coordinated across frequencies

Although rotational planes varied across grasp conditions, this variation was structured in three important ways. First, we found that the orientation of each rotation on a given condition was related to the orientation of the other rotation on the same condition, and to the overall rotational center (the “location”). To quantify this, we assessed how well the loadings (orientation) of one rotation / the location could predict those of another (Fig. 5a). Within each grasp condition, we used the loadings of the location or a rotation at one frequency to linearly predict those of a different frequency rotation, and quantified performance as the variance explained in the predicted rotation. Critically, the same prediction model was applied across all conditions. The location was predictive of rotation orientation for both M1 and F5 well above shuffle controls (Fig. 5b; second monkey in Supp. Fig. 6), though it was not perfectly predictive (M1: 74.8%, F5: 64.4% variance explained; p < 0.001). Prediction was statistically significant but poor in AIP (16.0%, p = 0.022, Wilcoxon signed-rank test), presumably because the activity related mostly to the visual cueing more than dynamic production of movement. Across all pairs of location and rotations, predictive performance was high in M1, F5, and AIP, explaining 90.5 ± 7.9%, 86.6 ± 11.4%, and 91.5 ± 8.1% of the variance, respectively (4-fold cross-validation; Fig. 5c, shown for M1; other areas and monkey in Supp. Fig. 6). When data were residualized to remove the common signal, predictive performance was lower but still above a shuffle control (Supp. Fig. 7). To determine whether this predictability exceeded that expected by chance, we compared prediction performance with a permutation-based null distribution generated by randomly reassigning component-condition relationships (10,000 shuffles). Cross-component predictions consistently exceeded the null distribution in all three cortical areas (permutation test, p<0.001 for all off-diagonal pairs; see prediction level above chance in Supp. Fig. 8; residualized in Supp. Fig. 9), demonstrating reliable relationships between different temporal components across conditions. These results suggest that although different frequencies occupy different subspaces of neural state space, their parameters are not independent and instead reflect a shared underlying structure.

**Figure 5:**
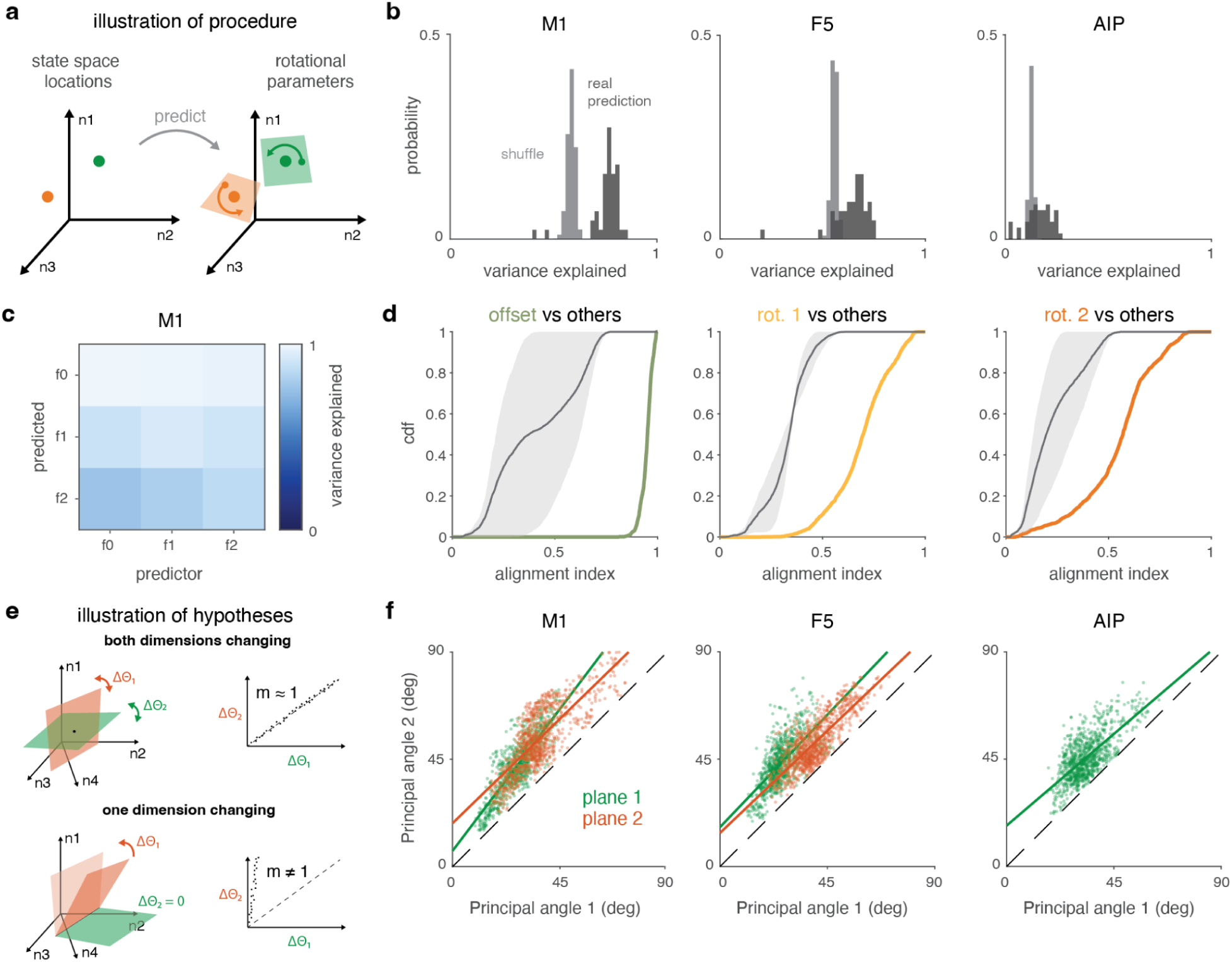
Reorientation of rotation subspaces is coordinated across frequencies and with state space location. **a.** Illustration of the LDR model, where the location of the neural population’s rotation center in state space predicts the orientation parameters of rotational dynamics. **b.** Fraction of population variance explained by predicting rotational parameters from state space location in M1, F5, and AIP (dark gray) compared to shuffled controls (light gray; p<0.001, permutation test with 10,000 shuffles). **c.** Mean variance explained in each rotation across conditions when predicted from a separate rotation in M1. Values along the diagonal are cross-validated and serve as an estimator of the performance ceiling due to noise. Off-diagonal predictions exceeded shuffled controls for all six comparisons (one-sided empirical p<0.001), whereas diagonal self-predictions did not differ from shuffled controls (p>0.05). **d.** Cumulative distribution function (CDF) of the alignment index for different-frequency rotational planes (gray) and same-frequency rotational planes (colored) across condition pairs in M1. Shaded gray regions indicate the minimum and maximum CDF values obtained across shuffles. **e.** Illustration of both dimensions of the rotation plane changing (top) vs. only one dimension of the rotation plane changing while the other is preserved, like a hinged door (bottom). **f.** Scatters of the two principal angles formed by each pair of corresponding-rotation planes. Each point corresponds to a condition pair; colors denote different rotation planes (different frequencies). Solid lines show linear fits; dashed identity line corresponds to both dimensions changing in proportion without noise. All correlations p < 0.001; slopes were 1.32 and 0.97 for M1 planes 1 and 2, 1.05 and 0.95 for F5 planes 1 and 2, and 0.77 for AIP plane 1.

Second, the subspaces used by different rotational frequencies were much more distinct than the subspace variation for a single frequency. Quantitatively, across all grasp-condition pairs, the subspace alignment was consistently higher for same-frequency than for different-frequency comparisons (Fig. 5d, M1 shown; other areas and monkey in Supp. Fig. 6; residualized in Supp. Fig. 7). This argues that a given subspace is selectively associated with a particular conserved temporal frequency rather than arising from arbitrary variation.

Finally, we found that it was the whole rotational subspace that changed across conditions, not just one dimension changing while the other remained fixed (Fig. 5e). To determine this, we considered pairs of conditions and asked whether the magnitude of change in one dimension predicted the magnitude of change in the other dimensions; i.e., whether the principal angles were correlated. Across all three cortical areas, principal angle changes were strongly correlated across rotational planes (Fig. 5f; second monkey in Supp. Fig. 6j; residualized in Supp. Fig. 7), using most of the possible range of angles and with the relationship having a slope of approximately 1. This structure was consistent with changes in both dimensions of the rotational plane being large or small together. Correlations were significant for both rotational-plane comparisons in M1 (Pearson’s r = 0.85 and 0.79, p < 0.001), both comparisons in F5 (r = 0.68 and 0.81, p < 0.001), and the single comparison in AIP (r = 0.70, p < 0.001).

### Geometry of dynamics captures behaviorally relevant information

The preceding analyses indicate that grasp-related population activity is organized around conserved temporal frequencies expressed through condition-dependent rotational subspaces. An important question is whether this geometry of dynamics captures behaviorally relevant information beyond what is available in representational models. One direct test of this idea is an encoding analysis. If grasp kinematics predict the geometry of the neural dynamics, and that geometry in turn reconstructs neural activity better than a direct encoding model, then the dynamical representation captures real structure that is not easily described as a simple representational mapping. Using encoding and decoding models of the loadings from LDR^10^, we tested whether grasp kinematics relate to the location and orientation of the rotations. For these analyses, we considered only M1 and F5, where LDR fits were stronger.

For each cortical area, we fit encoding models that used hand and finger kinematics to predict the condition-specific LDR parameters describing the orientation and magnitude of the rotational components (the loadings). Predicted LDR parameters were then used to reconstruct neural activity. We compared this approach with a linear encoding model that predicted neural activity directly from kinematics. Performance was assessed using leave-one-condition-out cross-validation and quantified as variance explained (equivalent to R^2^) for individual units. For both cortical areas, LDR predicted a large fraction of the neural variance (Fig. 6a; M1: 0.70; F5: 0.56), and modestly but significantly outperformed the linear model (p<0.001, Wilcoxon signed-rank test). This higher performance indicates that there is structure in the neural activity that is predicted from the geometry of the dynamics that is not predicted from a linear relationship with the kinematics directly. Predicted PETHs also approximately matched observed firing rates across grasp conditions (Fig. 6b), as expected, indicating that the low-dimensional LDR representation preserved much of the structure of neural responses throughout the grasp network. Results were similar for the second monkey (Fig. 6c; neural variance predicted: M1: 0.66; F5: 0.45; p<0.001 vs. linear direct model). To determine whether these conclusions depended on condition-specific mean responses, we repeated the analyses after residualizing the data. Encoding performance was still substantial and statistically significant for M1 of both monkeys and F5 of one monkey, but was substantially lower overall (Supp. Fig. 10a,b), with little difference between LDR and direct linear encoding. This argues that the condition-specific geometry of the dynamics relates appreciably to the condition-specific kinematics, with the lower performance presumably following from the low signal-to-noise ratio in residualized single-trial single-neuron data.

**Figure 6:**
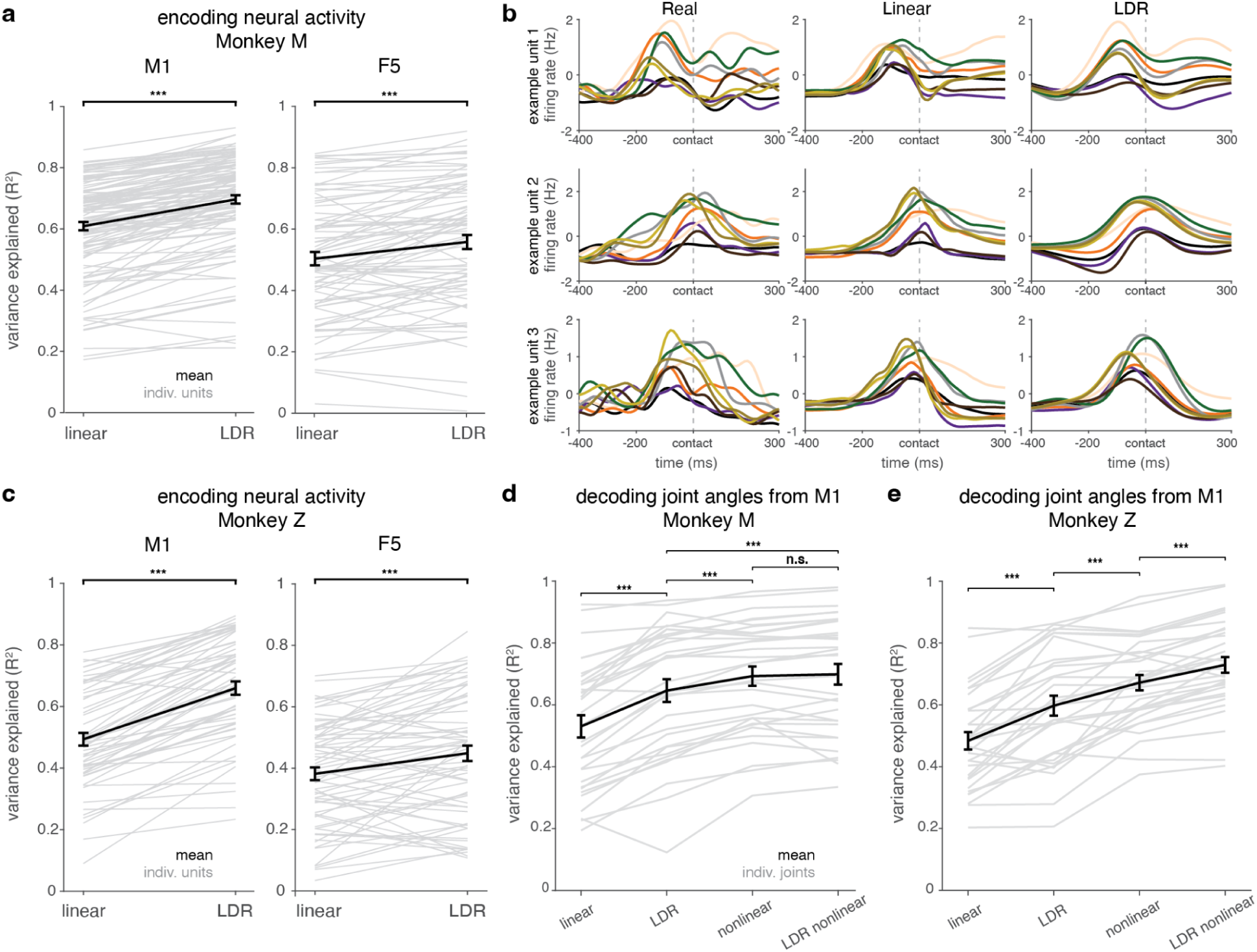
LDR geometry captures behaviorally relevant information. **a.** Encoding performance predicting neural activity from kinematics using direct linear regression (“linear”) and LDR with a linear relationship, evaluated using leave-one-condition-out cross-validation. Monkey M. Performance quantified as variance explained (R^2^). Gray lines indicate individual unit R^2^ values; black line indicates mean across units. AIP excluded for these analyses because of low variance explained by LDR. **b.** Example M1 neurons illustrating observed and model-predicted PETHs. Each row shows a different example neuron. **c.** Same as **a**, for monkey Z. **d.** Decoding performance for joint angle kinematics reconstructed from M1 activity using linear direct, LDR, nonlinear direct, and LDR nonlinear decoding models, evaluated using five-fold cross-validation over trials. Performance quantified as variance explained (R^2^). Gray lines indicate individual joints; black line indicates mean across joints. Monkey M. **e.** Same as **d**, for monkey Z.

We next fit decoding models, using the LDR parameters to predict kinematics. Because M1 is most directly linked to motor output, we focused on decoding hand and finger kinematics from M1 population activity alone. Joint angle trajectories were reconstructed using four approaches: direct linear regression, direct nonlinear regression (support vector regression with a nonlinear kernel), and LDR-based decoders with either linear or nonlinear regression. Performance was evaluated using five-fold cross-validation over trials and quantified as variance explained (R^2^) on the held-out data. As with encoding, kinematic decoding performance improved substantially and significantly using LDR vs. direct linear decoding (Fig. 6d,e; p<0.001, Wilcoxon signed-rank test). Using nonlinear regression either with LDR or with direct decoding further improved performance slightly but significantly above linear LDR (p<0.001 for both), and nonlinear LDR produced stronger decoding still for one monkey (p<0.001) but comparable performance to nonlinear direct decoding for the other. In the residualized data, decoding performance was again reduced for all methods, but LDR yielded a substantial improvement over direct decoding for both linear and nonlinear versions in both monkeys (Supp. Fig. 10c,d). This result supports a strong relationship of rotation location and orientation to kinematics. Overall, these results indicate that the geometric organization of the dynamics captures much of the behaviorally relevant nonlinear structure present in the neural population, enabling simple linear decoders to achieve performance approaching that of more flexible nonlinear models when sufficient task-related signal is present.

### Reach and grasp employ similar geometric strategies in different quantitative regimes for movement generation

Given that the angle between the common trajectory and the condition-specific activity was relatively small, and that the rotations varied through many dimensions but at smaller angles than in reach, we asked whether there might be a difference in the magnitude of how much the location differed across conditions for grasp vs. reach. To do so, we quantified two complementary measures (Fig. 7a). State space “location distance” measures the separation between the rotational centers of a pair of grasp conditions, whereas “rotation diameter” captures the maximum distance within a trajectory projected into that condition’s rotational plane (Methods).

**Figure 7:**
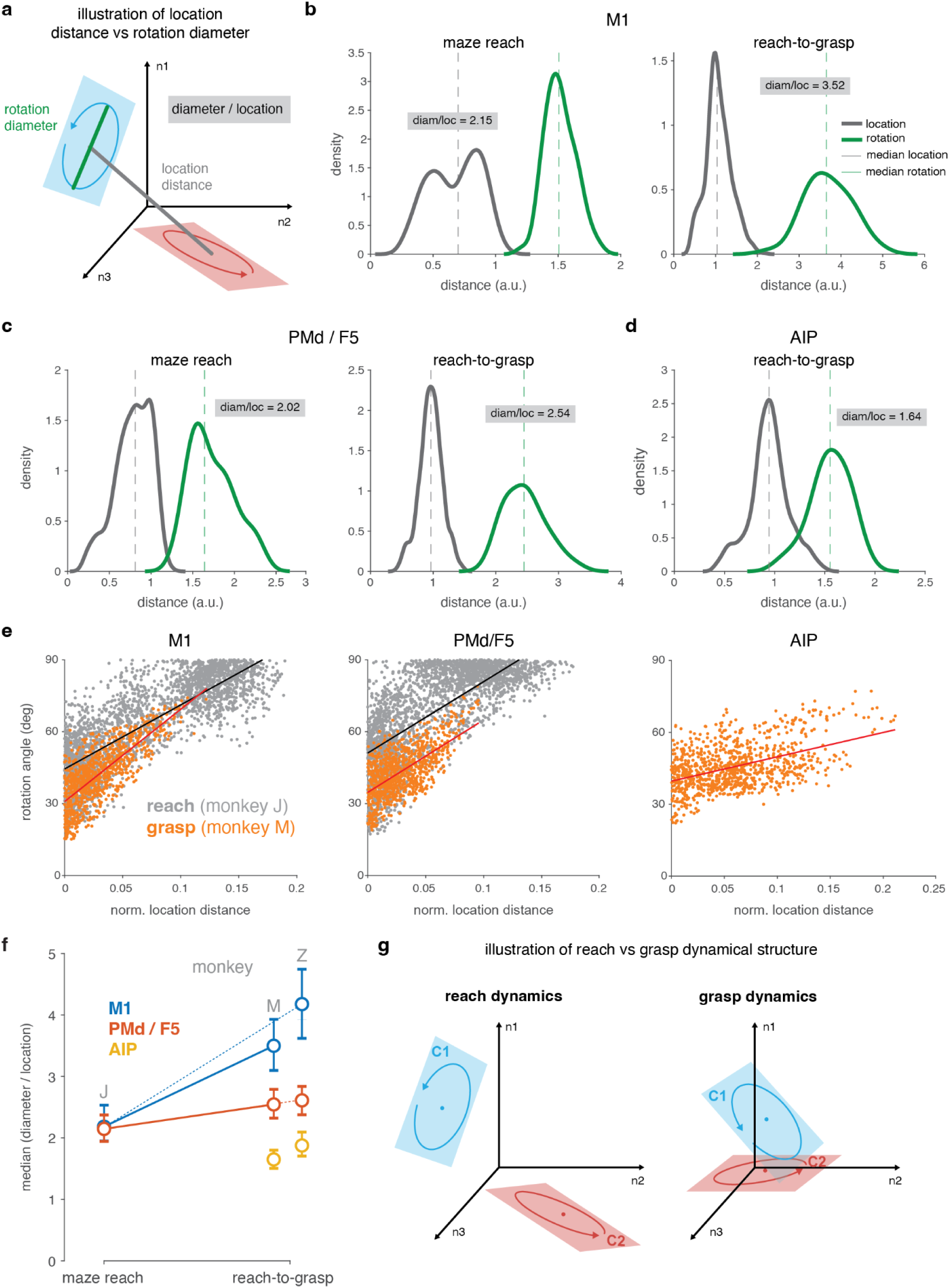
Grasp explores proportionately less of state space than reach. **a.** Illustration of the rotation-plane diameter and state space location distance metrics. Rotation-plane diameters are the scale of the rotational trajectories within a plane. Location distances are the distances between the condition-specific offsets (which form the rotational centers). **b.** Distribution of state space location distance across pairs of conditions (gray) and rotation-plane diameter (green) for the first oscillatory mode in M1 during the maze reach task (left) and reach-to-grasp task (right). **c.** Same as **b**, shown for PMd (maze reach task) or F5 (reach-to-grasp task). **d.** Same as **b**, shown for AIP during reach-to-grasp task. **e.** Angular difference in the rotational plane (first principal angle for lowest-frequency rotational plane) as a function of location distance. Points are pairs of conditions. Gray, reach data; orange, grasp data. Lines show linear regression fits. **f.** Summary of the diameter to location ratio across datasets. Error bars show 95% confidence intervals (bootstrap). **g.** Schematic contrasting dynamical structure during reach and grasp in neural state space.

Comparing the diameter of the rotations to the location separation, for M1, the rotations were relatively larger for grasp than for reach (64% larger for monkey M; Fig. 7b). For F5 (or PMd), this effect was somewhat smaller, with rotations only 25% larger for grasp than reach (Fig. 7c; second monkey in Supp. Fig. 11a). These effects were consistent across monkeys (Fig. 7f; monkey M, median ratio difference in M1: 1.29, 95% bootstrap CI [0.79, 1.79], p<0.0001; F5: 0.40, [0.08, 0.71], p = 0.013; monkey Z, M1: 1.98, [1.34, 2.60], p<0.0001; F5: 0.42, [0.11, 0.73], p = 0.01). These results suggest that the state space locations did not differ as much across conditions for grasp as for reach.

This observed difference, that rotations were larger relative to location distance for grasp than for reach, led to two further questions about the geometry of the dynamics in grasping. First, when considering two conditions, did the amount of change in the rotational plane scale with difference in location? The answer was yes, at least for M1 and F5: there was a strong relationship between the location distance for a pair of rotations and the angular difference between their rotational planes (Fig. 7e, left and center; orange points). For AIP, the relationship was present but expectedly much weaker (Fig. 7e, right).

Second, given that grasp-related activity exhibited less variation in rotation plane angle (Fig. 4i) than observed for reach^10^, we asked: were the smaller angles for grasp due to smaller location distances between conditions, or a different relationship between location and orientation? In M1, we found that the smaller distances between state space locations explained most of the difference (Fig. 7e, left; orange points generally have smaller location values than gray points but a generally similar fit; second monkey in Supp. Fig. 11b). For F5, the effect was mixed: location distances were still generally smaller for grasp than reach, but the same location distance also led to less rotation angle difference for grasp than reach (Fig. 7e, center; orange line vertically offset from gray line). These results together suggest that the smaller variation in rotational orientation for grasp than reach reflects use of a smaller range of location (Fig. 7g). Analyzing the residualized data confirmed this relationship, showing similar fits for the rotation-angle vs. location-difference relationship for reach and grasp (Supp. Fig. 12). This residual analysis suggests that the mild disagreement between reach and grasp in the full data can be explained by the inclusion of the unchanging, non-orthogonal common signal proportionally reducing the rotational plane differences across conditions.

## Acknowledgements

The authors thank H. Scherberger and S. Schaffelhofer for sharing their data, D. Sabatini, A. Pavuluri, S. Bensmaia, L. Okorokova, A. Sobinov, and J. Downey for discussions of this work, M. Greaney for the monkey illustration, and H. Scherberger for critical review of the manuscript. This work was funded by The Computational Neuroscience Training Grant (ZG), NIH NINDS R01 NS125270 (MK), the NSF-Simons National Institute for Theory and Mathematics in Biology via grants NSF DMS-2235451 and Simons Foundation MP-TMPS-00005320 (MK), and The University of Chicago.

## Author contributions

Both authors contributed to all parts of conceptualization, interpretation, writing, and editing. Z.G. performed the analyses with assistance from M.K. M.K. raised the funding for the work.

## SUPPLEMENTARY FIGURES

**Supplementary figure 1:**
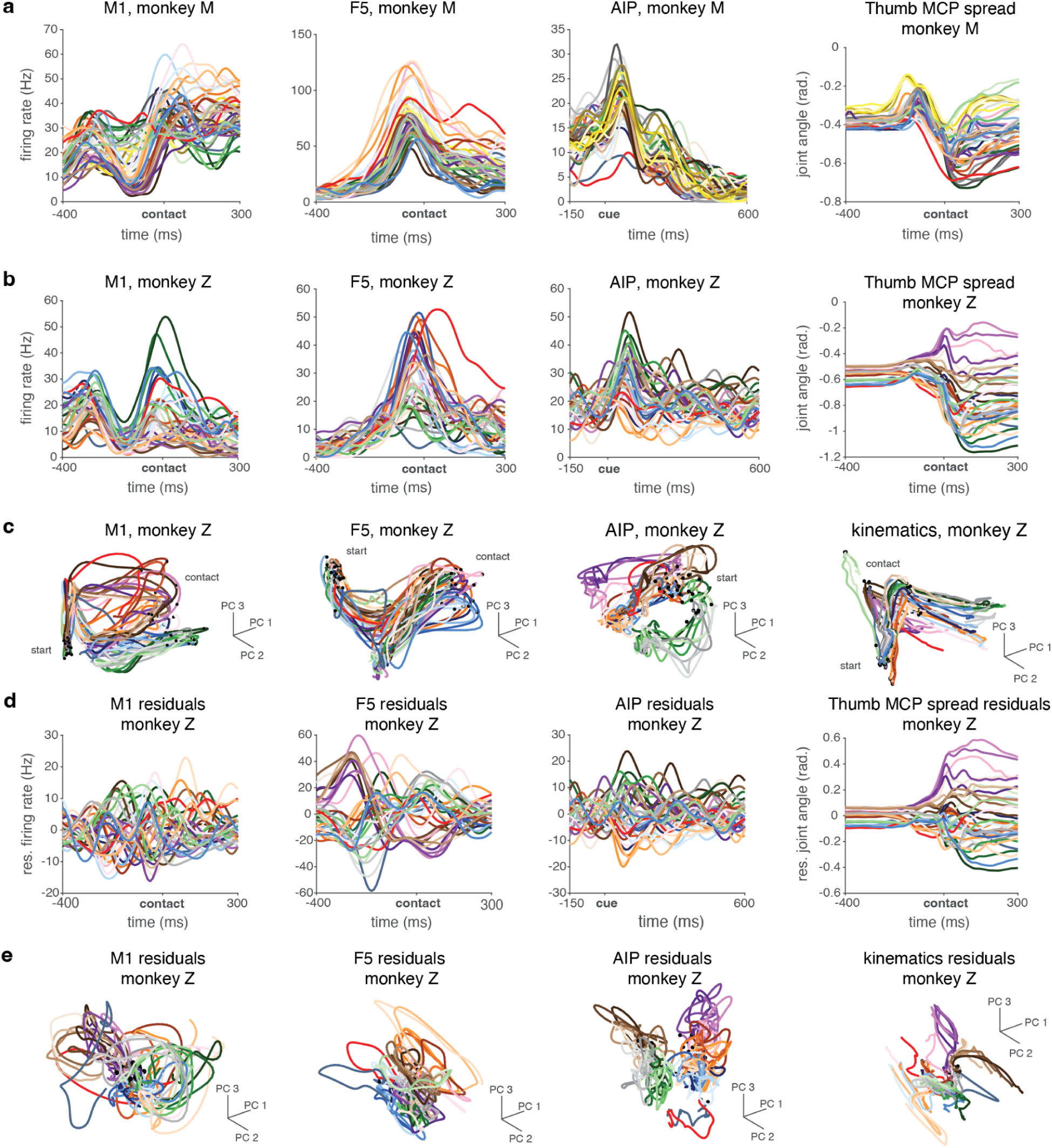
Additional neural and kinematic responses in the reach-to-grasp task. **a.** Example PETHs across the grasping network for monkey M (first three panels), aligned to contact for M1 and F5 and cue for AIP; and an example joint time course (last panel). **b.** Same as **a** for monkey Z. **c.** PCA trajectories of population activity in M1, F5, and AIP neural state space, and joint angle kinematics for monkey Z, colored by grasp condition. **d.** PETHs for example units (first three panels) and joint (last panel) after subtracting the condition-mean for monkey Z. Different units than shown in **b**. **e.** PCA trajectories of neural population activity (M1, F5, AIP) and joint angle kinematics after subtraction of the condition-mean for monkey Z, colored by grasp condition.

**Supplementary figure 2:**
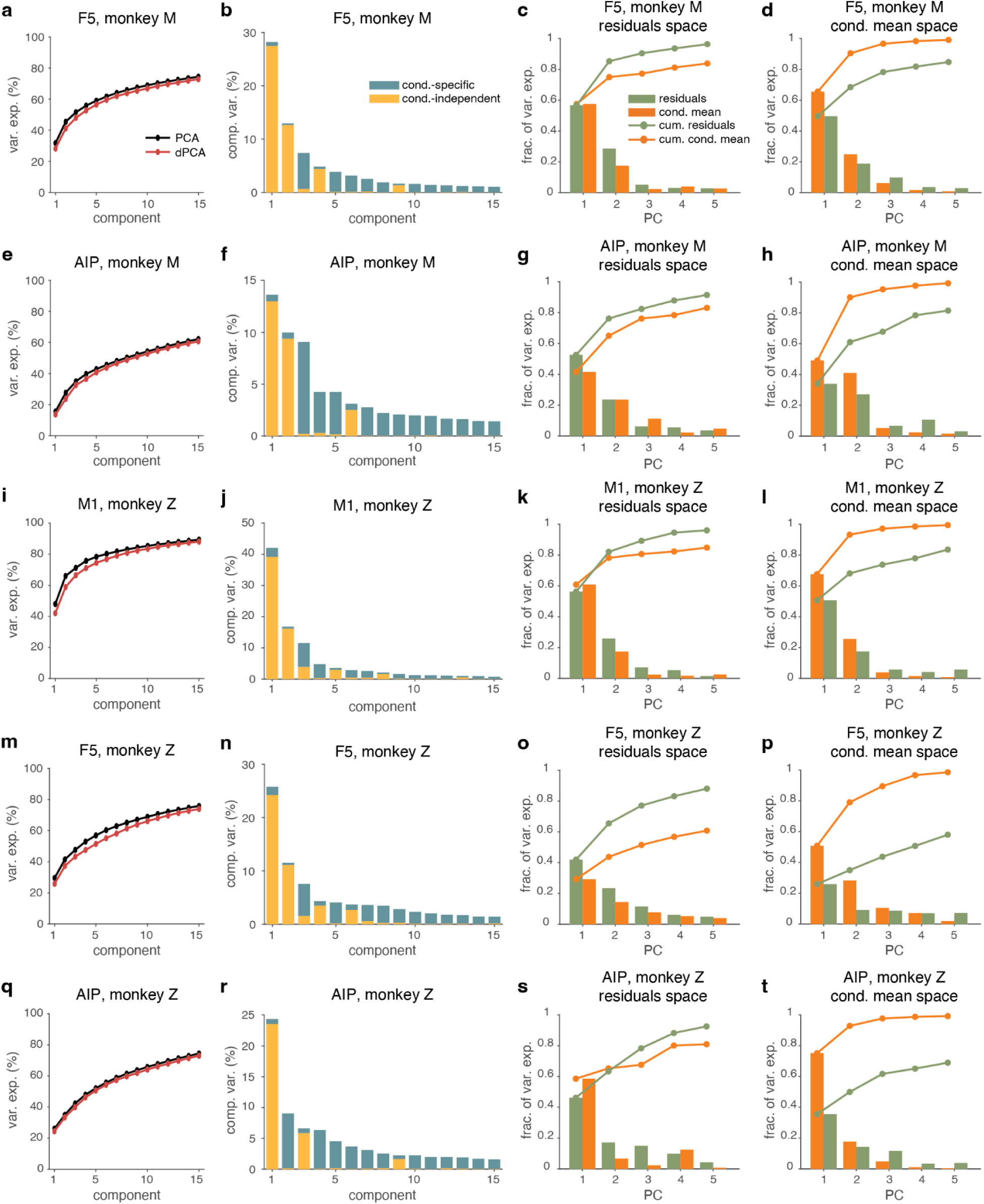
Overlap in condition-independent and grasp condition-specific activity, additional data. **a.** Cumulative variance explained by PCA (black) and dPCA (red) for F5 population activity in monkey M. **b.** Variance explained by individual demixed principal components in F5 for monkey M. Bars indicate the proportion of total variance explained and are partitioned into condition-specific variance (blue) and condition-independent variance (yellow). Note that the basis is non-orthogonal and so the variance contained in the condition-independent and the condition-specific subspaces is partly shared; the total variance explained by a component may therefore be less than the sum of its marginalized parts. **c.** Variance explained of the condition-mean (orange) and of the residual activity (green) in residuals space in F5 for monkey M. **d.** Variance explained of the condition-mean and of the residual activity in condition-mean space in F5 for monkey M. **e-h.** Same as **a-d** in AIP for monkey M. **i-l.** Same as **a-d** in M1 for monkey Z. **m-p.** Same as **a-d** in F5 for monkey Z. **q-t.** Same as **a-d** in AIP for monkey Z.

**Supplementary figure 3:**
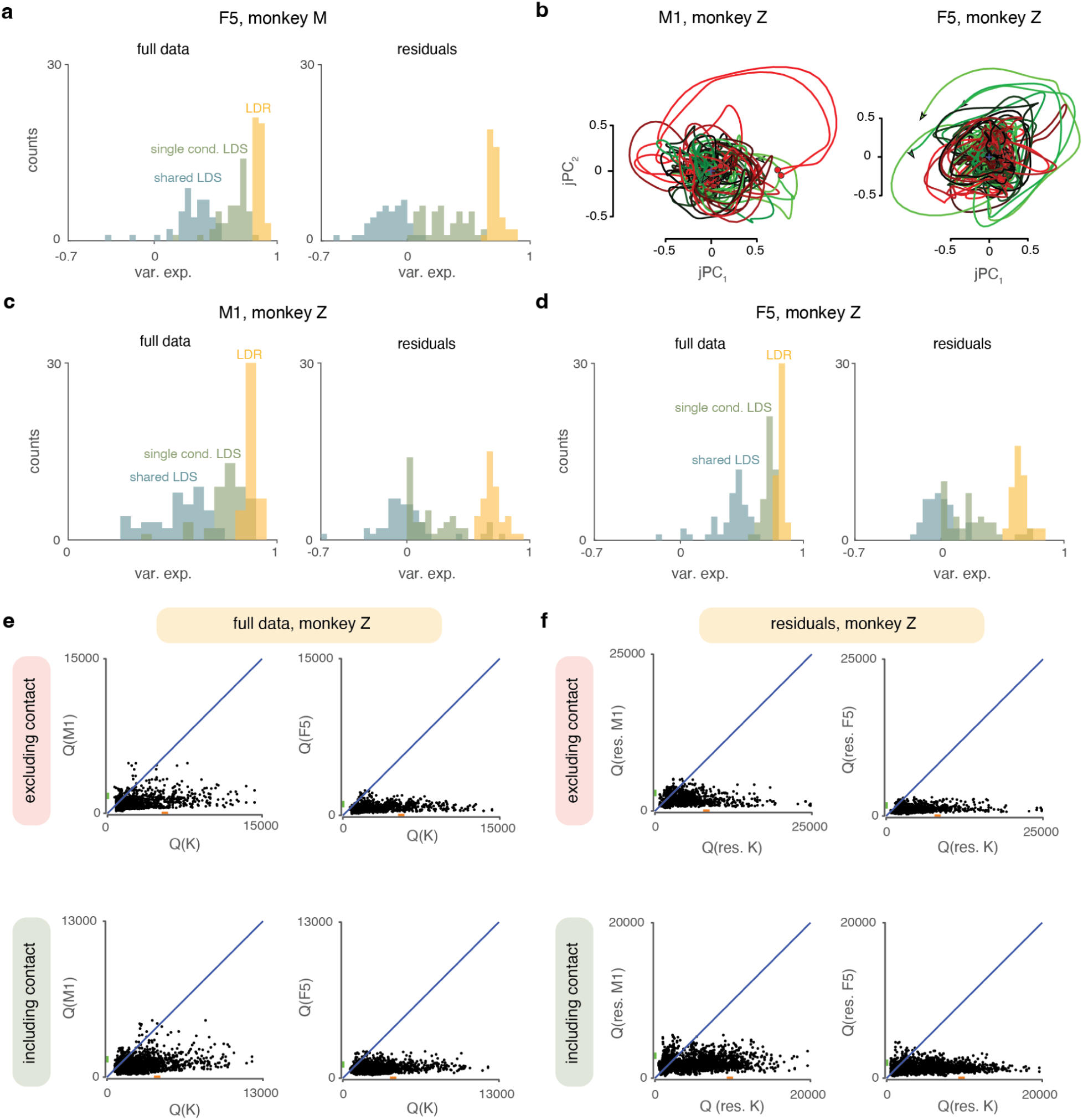
Testing dynamical systems properties in grasp-related neural activity, additional data. **a.** Cross-validated population variance explained by a shared LDS fit across conditions (blue), condition-specific LDSs fit separately to each grasp condition (green), and the LDR model (yellow). Left, fits to the full data; right, fits after subtracting the condition-mean. **b.** jPCA projections of trial-averaged neural population activity for individual grasp conditions in M1 and F5 for monkey Z. **c.** Same as **a** in M1 for monkey Z. **d.** Same as **a** in F5 for monkey Z. **e.** Trajectory tangling of full neural activity in M1 and F5 (Q(M1), Q(F5)) plotted against kinematic tangling (Q(K)) for analyses including only the pre-contact period (top) or including the full trial including object hold time (bottom) for monkey Z. **f.** Same as **e**, but computed using residualized neural activity (Q(res. M1), Q(res. F5)) and residualized kinematics (Q(res. K)) by subtracting the condition-mean.

**Supplementary figure 4:**
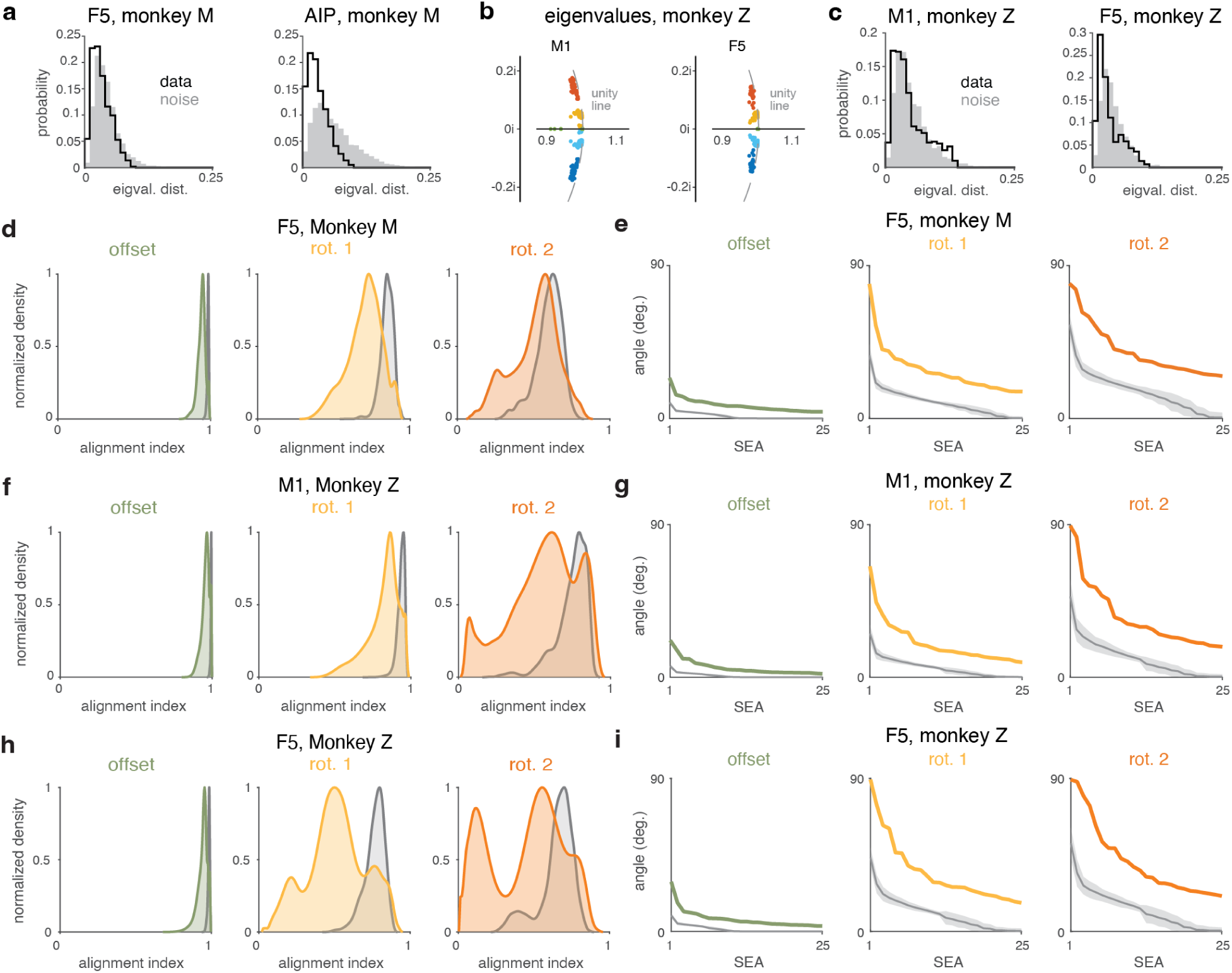
Conserved temporal frequencies are expressed through condition-specific geometry, additional data. **a.** Eigenvalue distance between conditions for F5 and AIP of monkey M, computed as the mean distance between corresponding (closest-frequency) eigenvalues in the complex plane. Noise distribution computed by partitioning the set of trials into halves and computing eigenvalues from the two halves for the same condition separately. **b.** Eigenvalues of the inferred dynamics for all grasp conditions in M1 and F5 for monkey Z. **c.** Same as **a** for M1 and F5 of monkey Z. **d.** Alignment indices between corresponding (same-frequency) rotational planes across pairs of grasp conditions (colored), compared with the distribution expected from estimation noise (gray) in F5 for monkey M. Distribution shown with kernel density estimation. **e.** Subspace excursion angles between corresponding rotational planes across conditions (colored), along with angles expected from estimation noise (gray) in F5 for monkey M. For the noise distribution, the line indicates the mean; shaded regions indicate ±1 s.d. **f.** Same as **d** in M1 for monkey Z. **g.** Same as **e** in M1 for monkey Z. **h.** Same as **d** in F5 for monkey Z. **i.** Same as **e** in F5 for monkey Z.

**Supplementary figure 5:**
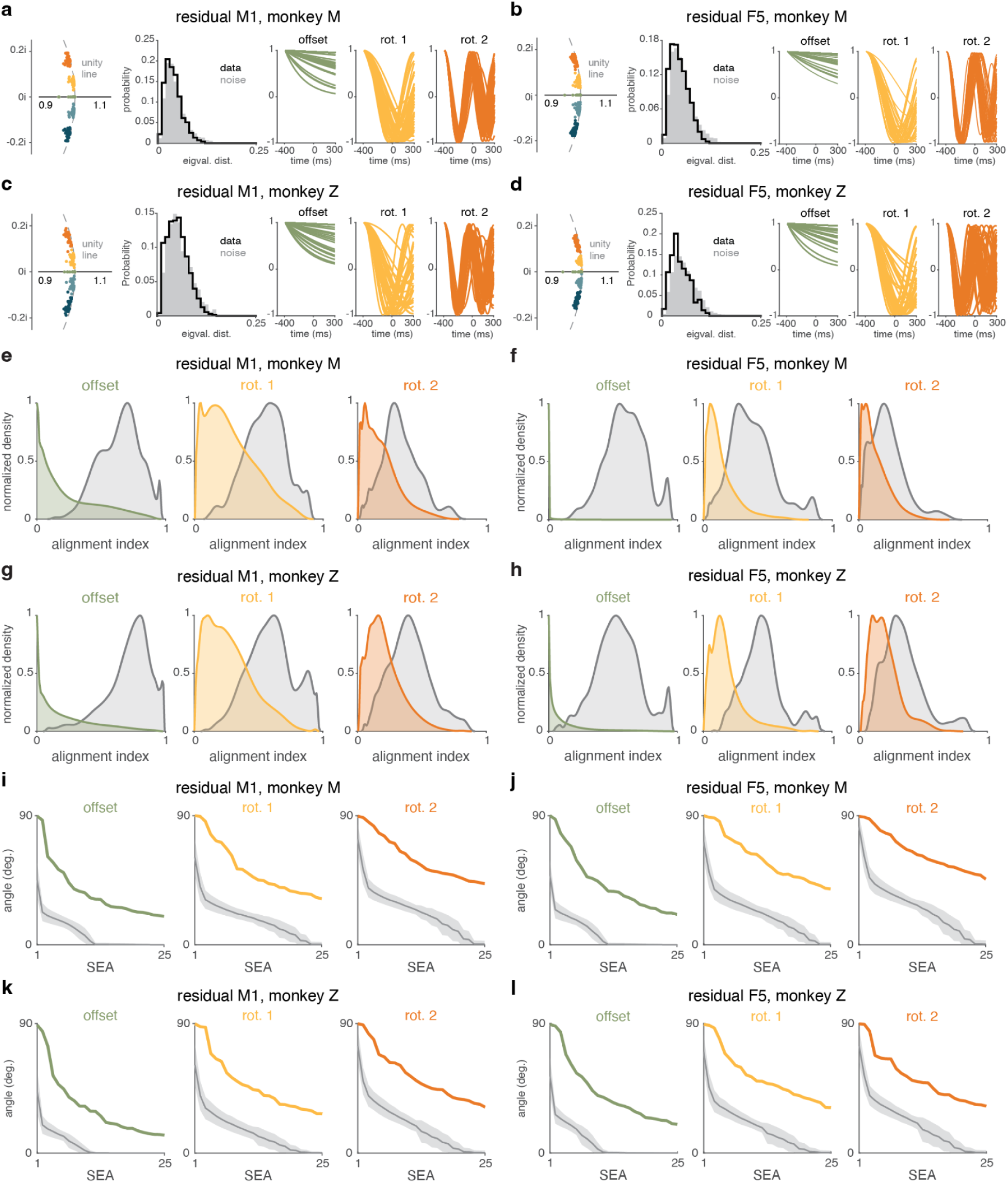
Conserved temporal frequencies are expressed through condition-specific geometry in residualized data. **a.** Eigenvalues of the inferred dynamics for all grasp conditions in M1 after residualization (left), eigenvalue distance between conditions in M1 after residualization compared with expected distribution due to noise (second panel), recovered temporal basis functions across all conditions in M1 after residualization aligned to contact (right three panels). Object hold epoch included. Monkey M. **b.** Same as **a** in F5 after residualization for monkey M. **c.** Same as **a** in M1 after residualization for monkey Z. **d.** Same as **a** in F5 after residualization for monkey Z. **e.** Alignment indices between corresponding (same-frequency) rotational planes across pairs of grasp conditions (colored), compared with the distribution expected from estimation noise (gray) in M1 after residualization. Distribution shown with kernel density estimation. Monkey M. **f.** Same as **e** for F5 after residualization for monkey M. **g.** Same as **e** for M1 after residualization for monkey Z. **h.** Same as **e** for F5 after residualization for monkey Z. **i.** Subspace excursion angles between corresponding rotational planes across conditions (colored), along with angles expected from estimation noise (gray) in M1 after residualization for monkey M. For the noise distribution, the line indicates the mean; shaded regions indicate ±1 s.d. **j.** Same as **i** for F5 after residualization for monkey M. **k.** Same as **i** for M1 after residualization for monkey Z. **l.** Same as **i** for F5 after residualization for monkey Z.

**Supplementary figure 6:**
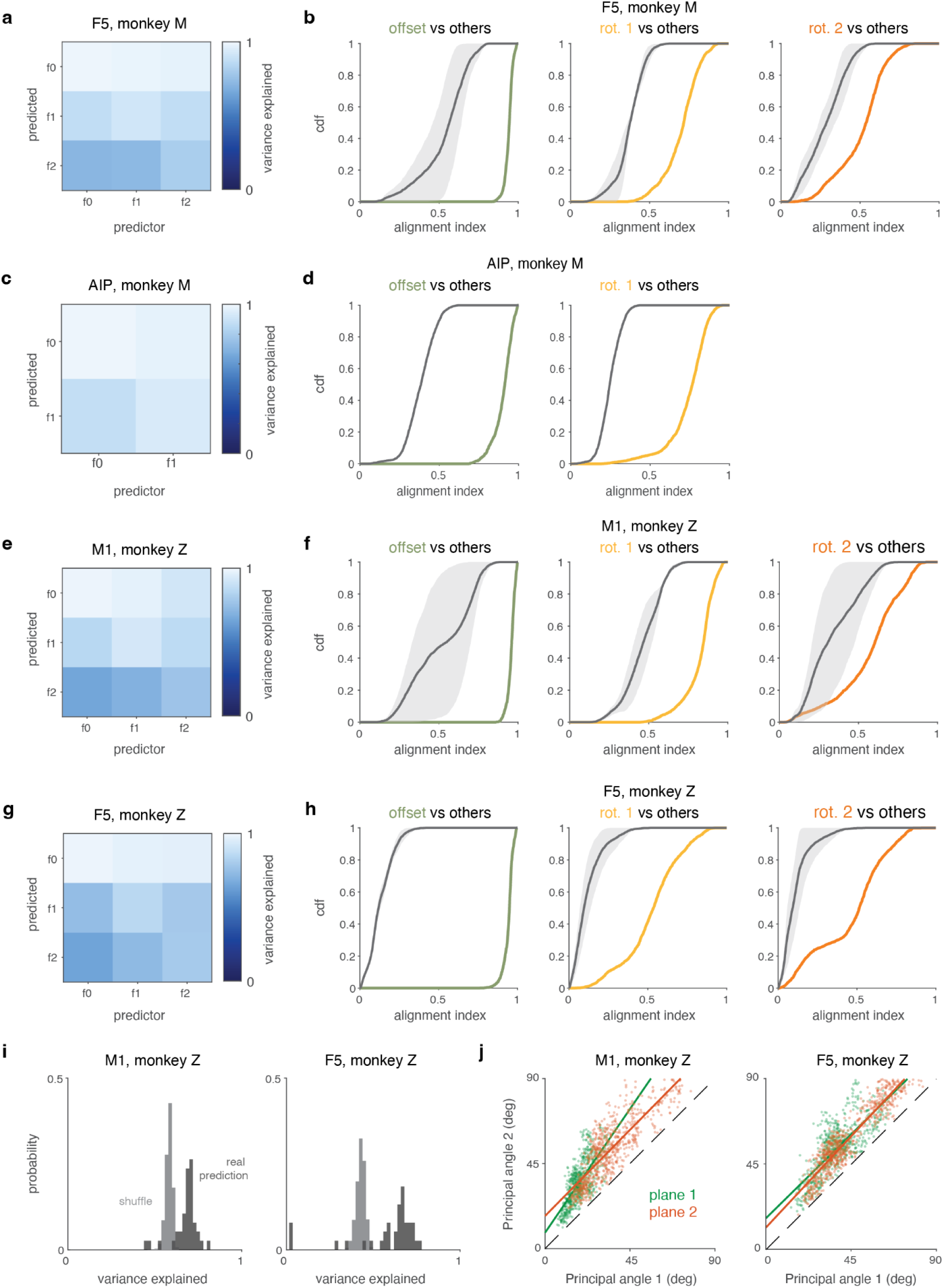
Reorientation of rotation subspaces is coordinated across frequencies and with state space location, additional data. **a.** Mean variance explained in each rotation across conditions when predicted from a separate rotation in F5 for monkey M. Values along the diagonal are cross-validated and serve as an estimator of the performance ceiling due to noise. Significance was assessed using a shuffle test (10,000 shuffles) in which component-condition relationships were randomly reassigned before model fitting. Off diagonal predictions exceeded shuffled controls (empirical one-sided p<0.001 for all comparisons), whereas diagonal self-predictions did not differ from shuffle controls (p>0.05). **b.** Cumulative distribution function of the alignment index for different-frequency rotational planes (gray) and same-frequency rotational planes (colored) across condition pairs in F5 for monkey M. Shaded gray regions indicate the minimum and maximum CDF values obtained across shuffles. **c.** Same as **a** in AIP for monkey M. Off-diagonal predictions exceeded shuffled controls (empirical one-sided p<0.001), whereas diagonal self-predictions did not differ from shuffle controls (p>0.05). **d.** Same as **b** for AIP in monkey M. **e.** Same as **a** in M1 for monkey Z. Off-diagonal predictions exceeded shuffled controls for 4 of 6 component pairs (one-sided empirical shuffle test, 10,000 shuffles, p<0.05), whereas diagonal self-predictions did not differ from shuffle controls (p>0.05). **f.** Same as **b** in M1 for monkey Z. **g.** Same as **a** in F5 for monkey Z. Off-diagonal predictions exceeded shuffled controls for 5 of 6 component pairs (one-sided empirical shuffle test, 10,000 shuffles, p<0.05), whereas diagonal self-predictions did not differ from shuffle controls (p>0.05). **h.** Same as **b** in F5 for monkey Z. **i.** Fraction of population variance explained by predicting rotational parameters from state space location in M1 (left) and F5 (right; dark gray) compared to shuffled controls (light gray) for monkey Z (p<0.001, permutation test with 10,000 shuffles). **j.** Scatters of the two principal angles formed by each pair of corresponding-rotation planes in M1 and F5 for monkey Z. Each point corresponds to a condition pair; colors denote different rotation planes (different frequencies). Solid lines show linear fits; dashed identity line corresponds to both dimensions changing in proportion without noise. All correlations p<0.001; slopes were 1.46 and 1.01 for M1 planes 1 and 2, and 0.98 and 1.07 for F5 planes 1 and 2.

**Supplementary figure 7:**
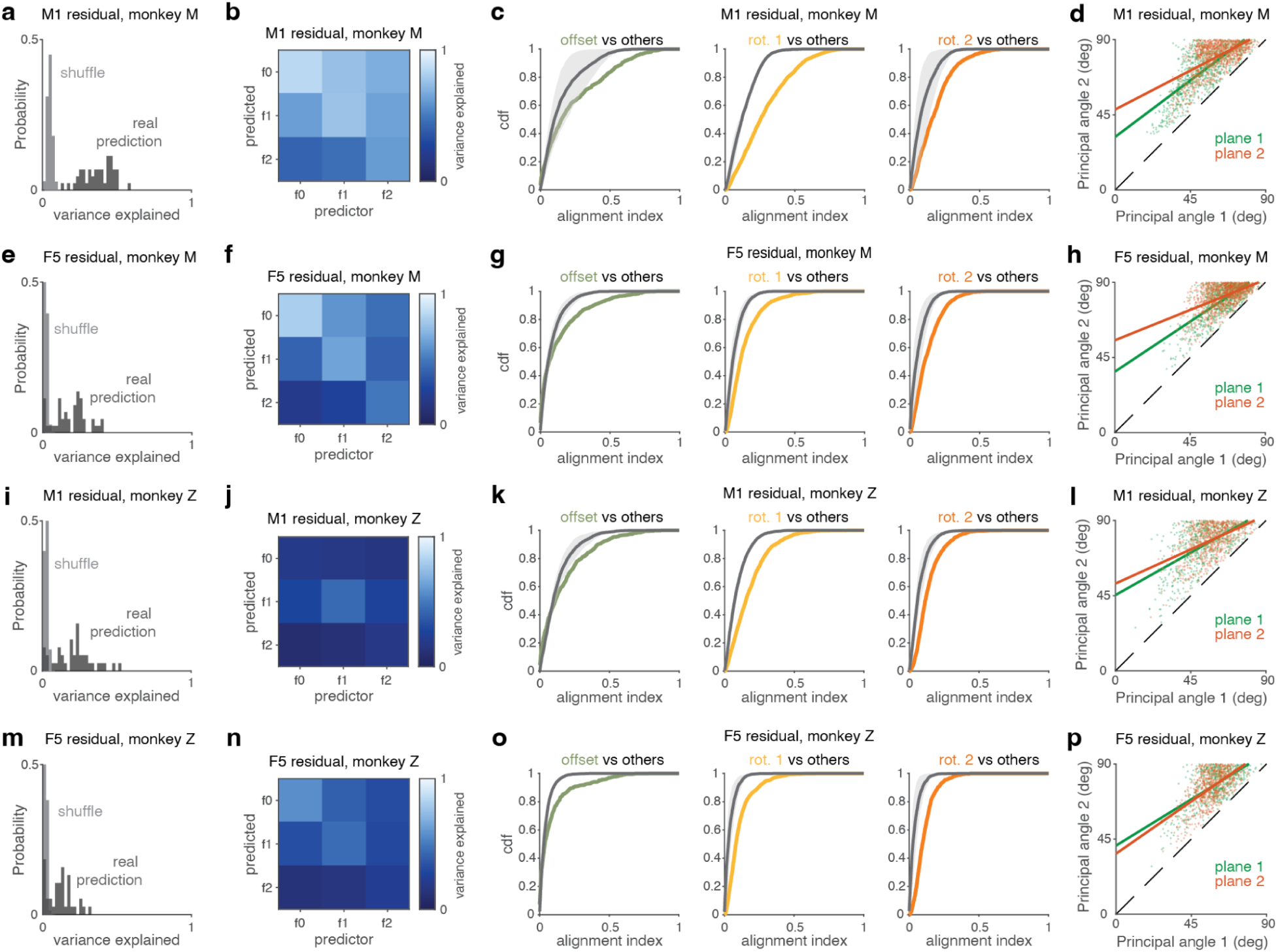
Reorientation of rotation subspaces is coordinated across frequencies and with state space location, residualized data. **a.** Fraction of population variance explained by predicting rotational parameters from state space location in M1 residual and F5 residual (dark gray) compared to shuffled controls (light gray) for monkey M (p<0.001, permutation test with 10,000 shuffles). **b.** Mean variance explained in each rotation across conditions when predicted from a separate rotation in M1 residual for monkey M. Values along the diagonal are cross-validated and serve as an estimator of the performance ceiling due to noise. Significance was assessed using a shuffle test (10,000 shuffles) in which component-condition relationships were randomly reassigned before model fitting. Off diagonal predictions exceeded shuffled controls (empirical one-sided p<0.001 for all comparisons), whereas diagonal self-predictions did not differ from shuffle controls (p>0.05). **c.** Cumulative distribution function of the alignment index for different-frequency rotational planes (gray) and same-frequency rotational planes (colored) across condition pairs in M1 residual for monkey M. Shaded gray regions indicate the minimum and maximum CDF values obtained across shuffles. **d.** Scatters of the two principal angles formed by each pair of corresponding-rotation planes in M1 residual for monkey M. Each point corresponds to a condition pair; colors denote different rotation planes (different frequencies). Solid lines show linear fits; dashed identity line corresponds to perfect pivoting without noise. All correlations p<0.001. **e.** Same as in **a** in F5 residuals for monkey M (p<0.001, permutation test with 10,000 shuffles). **f.** Same as **b** in F5 residuals for monkey M. Off-diagonal predictions exceeded shuffled controls (empirical one-sided p<0.001), whereas diagonal self-predictions did not differ from shuffle controls (p>0.05). **g.** Same as **c** for F5 residuals in monkey M. **h.** Same as in **d** in F5 residuals for monkey M. All correlations p<0.001. **i.** Same as in **a** in M1 residuals for monkey Z (p<0.001, permutation test with 10,000 shuffles). **j.** Same as **b** in M1 residuals for monkey Z. Off-diagonal predictions exceeded shuffled controls (one-sided empirical shuffle test, 10,000 shuffles, p<0.05), whereas diagonal self-predictions did not differ from shuffle controls (p>0.05). **k.** Same as **c** in M1 residuals for monkey Z. **l.** Same as in **d** in M1 residuals for monkey Z. All correlations p<0.001. **m.** Same as in **a** in F5 residuals for monkey Z (p<0.001, permutation test with 10,000 shuffles). **n.** Same as **b** in F5 residuals for monkey Z. Off-diagonal predictions exceeded shuffled controls (one-sided empirical shuffle test, 10,000 shuffles, p<0.05), whereas diagonal self-predictions did not differ from shuffle controls (p>0.05). **o.** Same as **c** in F5 residuals for monkey Z. **p.** Same as in **d** in F5 residuals for monkey Z. All correlations p<0.001.

**Supplementary figure 8:**
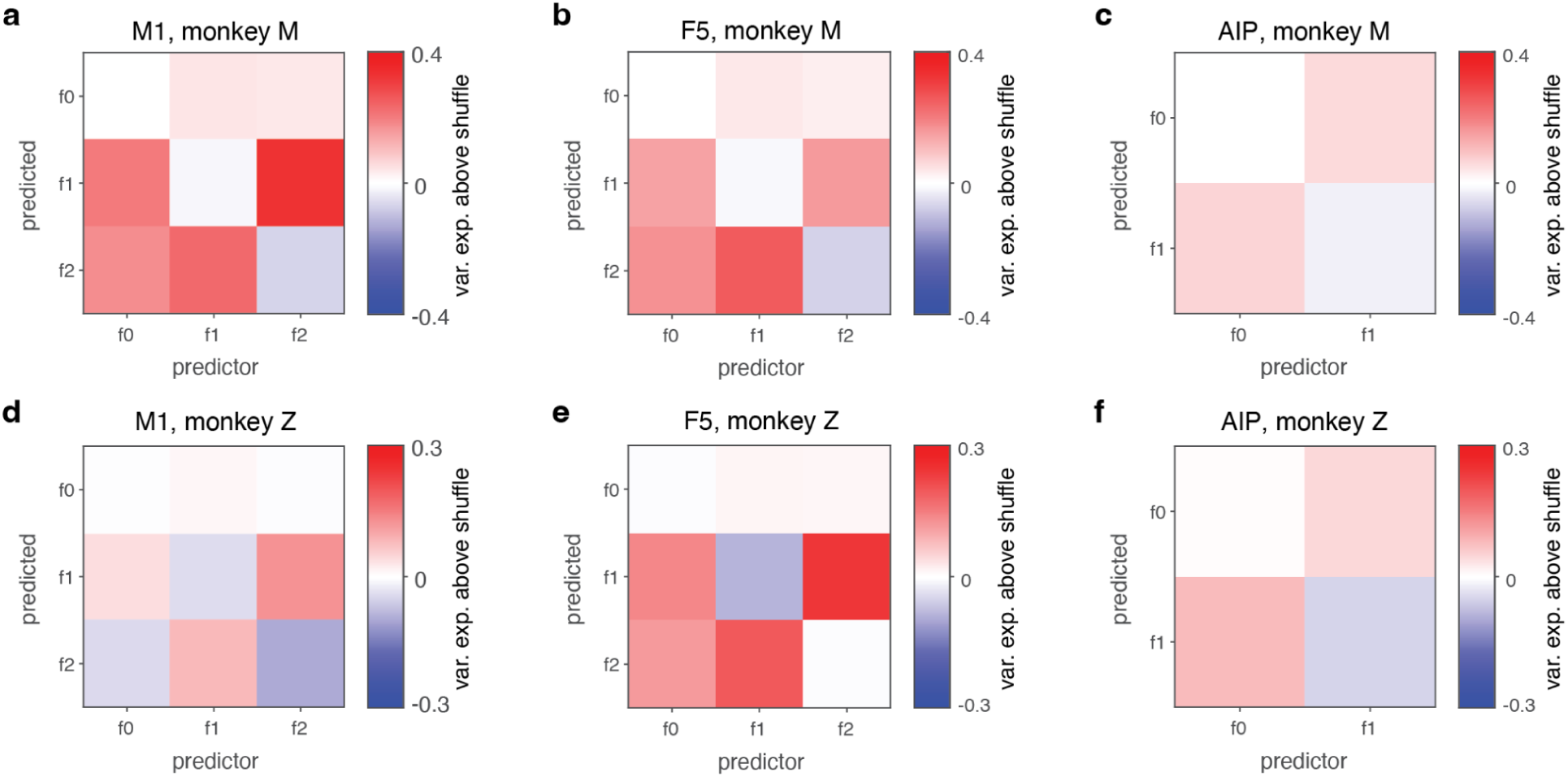
Prediction of temporal basis function components beyond condition-shuffled control. **a.** Heatmap of cross-validated variance explained between temporal components above a condition-shuffled control in M1 for monkey M. **b.** Same as **a** in F5 for monkey M. **c.** Same as **a** in AIP for monkey M. **d.** Same as **a** in M1 for monkey Z. **e.** Same as **a** in F5 for monkey Z. **f.** Same as **a** in AIP for monkey Z.

**Supplementary figure 9:**
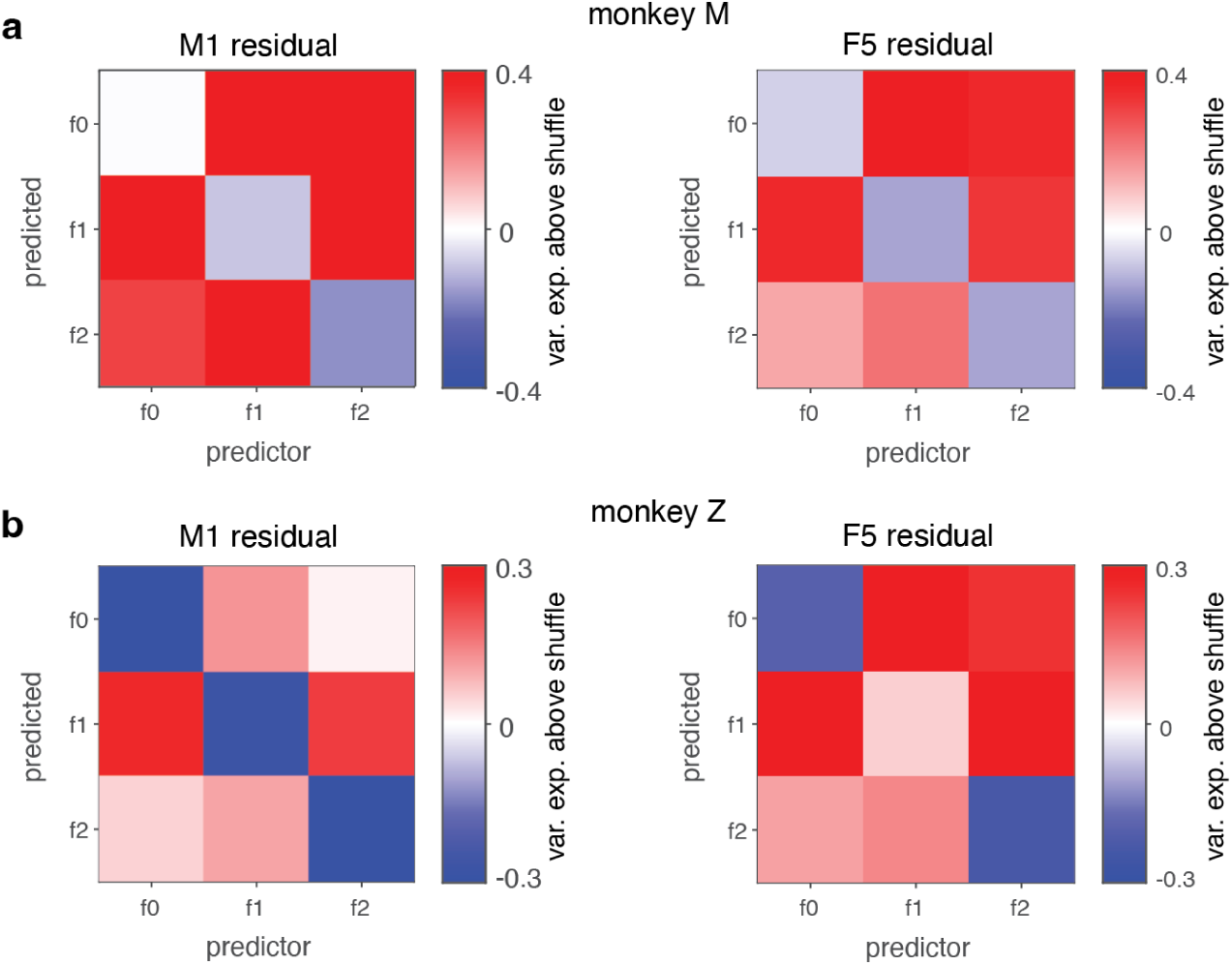
Prediction of temporal basis function components beyond condition-shuffled control, residualized data. **a.** Heatmap of cross-validated variance explained between temporal components above a condition-shuffled control in M1 residual (left) and F5 residual (right) for monkey M. **b.** Same as **a** for monkey Z.

**Supplementary Figure 10:**
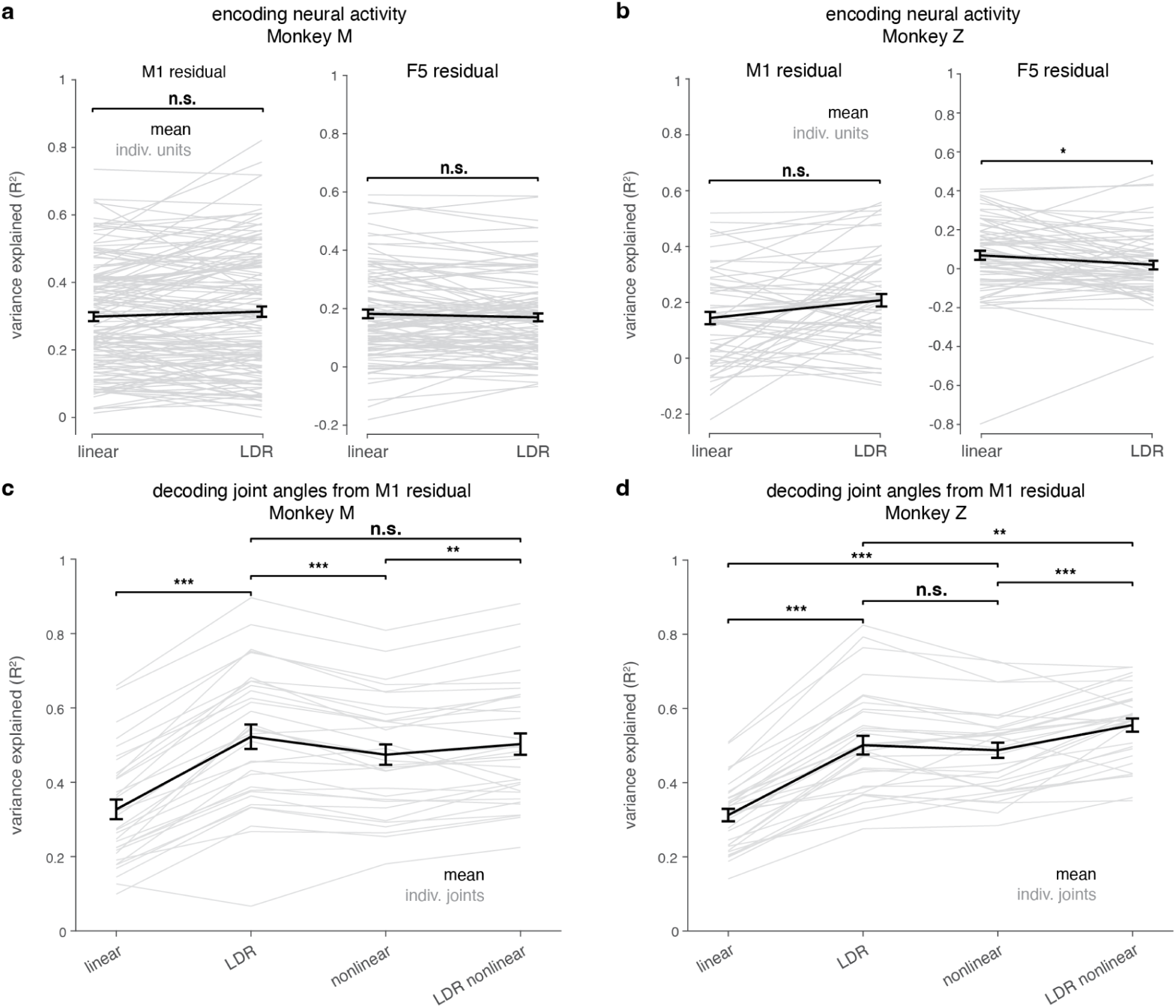
LDR geometry captures behaviorally relevant information, residualized data. **a.** Encoding performance predicting neural activity from kinematics using direct linear regression (“linear”) and LDR with a linear relationship, evaluated using leave-one-condition-out cross-validation. Monkey M. Performance quantified as variance explained (R^2^). Gray lines indicate individual unit R^2^ values; black line indicates mean across units. AIP excluded for these analyses because of low variance explained by LDR. **b.** Same as **a**, for monkey Z. **c.** Decoding performance for joint angle kinematic residuals reconstructed from M1 activity using linear, LDR, nonlinear, and LDR nonlinear decoding models, evaluated using five-fold cross-validation over trials. Performance quantified as variance explained (R^2^). Gray lines indicate individual joints; black line indicates mean across joints. Statistical comparisons were performed between adjacent decoding models. Monkey M. **d.** Same as **c**, for monkey Z.

**Supplementary figure 11:**
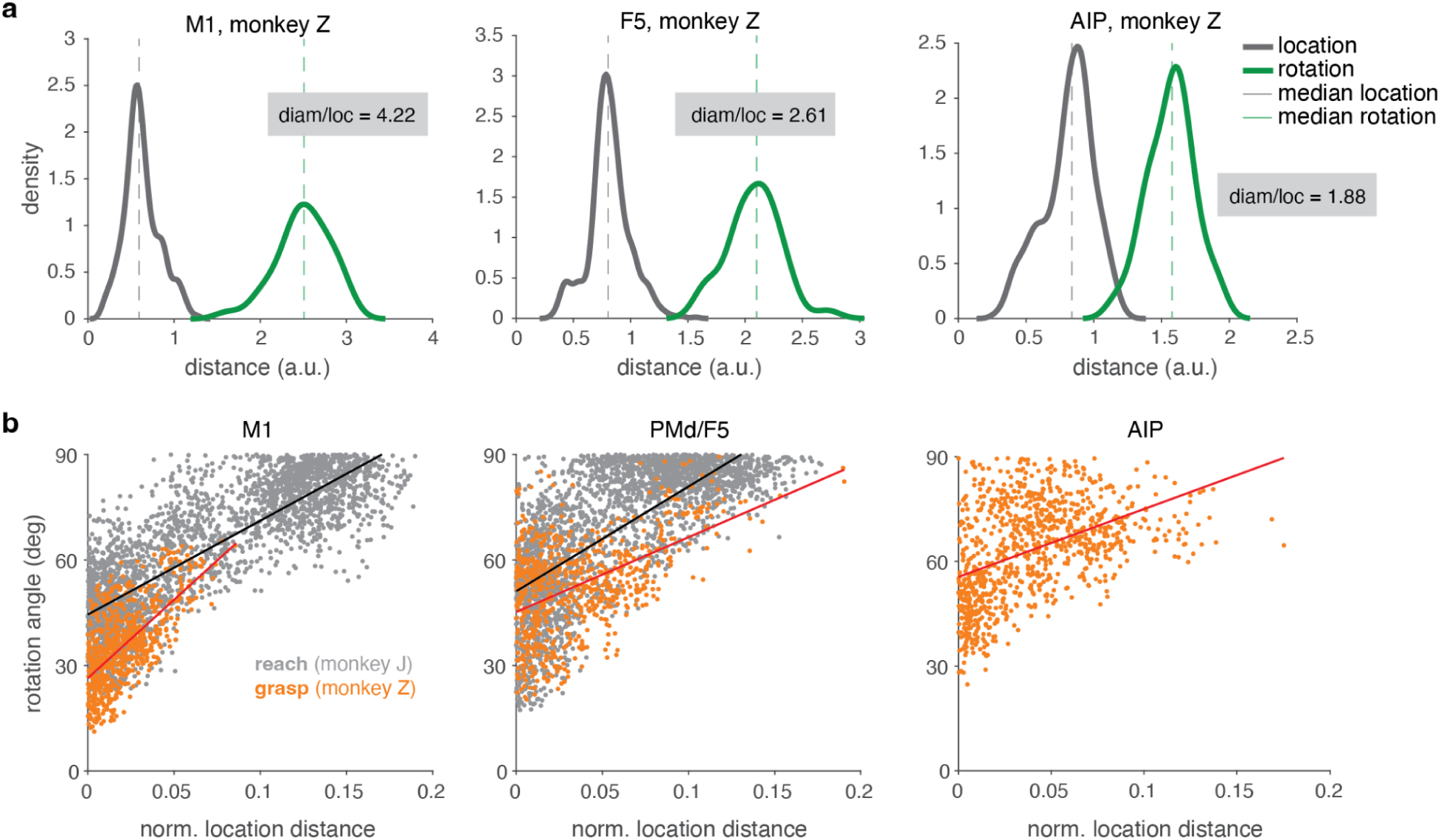
Grasp explores proportionately less of state space than reach, additional data. **a.** Distribution of state space location distance across pairs of conditions (gray) and rotation-plane diameter (green) for the first oscillatory mode during the reach-to-grasp task in M1 (left), F5 (center), and AIP (right) for monkey Z. **b.** Angular difference in the rotational plane (first principal angle for lowest-frequency rotational plane) as a function of location distance in M1 (left), PMd/F5 (center), and AIP (right) for monkey Z. Points are pairs of conditions. Gray, reach data; orange, grasp data. Lines show linear regression fits.

**Supplementary figure 12:**
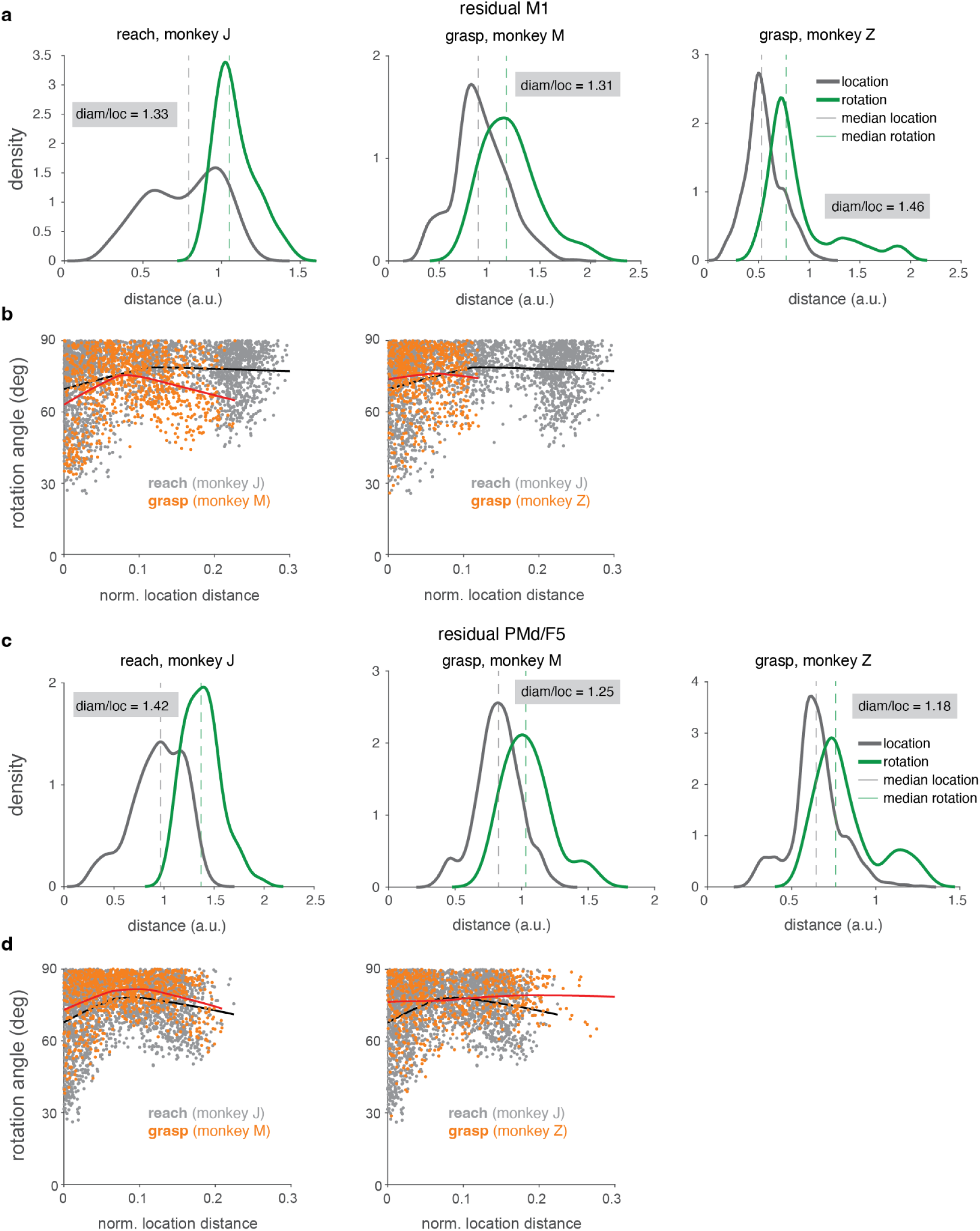
Relationship of location distance to rotation angle is more similar for reach and grasp in residualized data. **a.** Distribution of state space location distance across pairs of conditions (gray) and rotation-plane diameter (green) for the first oscillatory mode during maze reach (left) and reach-to-grasp task for monkey M (center) and reach-to-grasp task for monkey Z (right) in M1 after residualization. **b.** Angular difference in the rotational plane (first principal angle for lowest-frequency rotational plane) as a function of location distance in comparison to reach for monkey M (left) and monkey Z (right) in M1 after residualization. Points are pairs of conditions. Gray, reach data; orange, grasp data. Lines show LOWESS fits. **c.** Same as **a** in PMd/F5 after residualization. **d.** Same as **b** in PMd/F5 after residualization.

## DISCUSSION

The dynamics underlying the generation of grasp commands^39^ – and indeed whether the cortical grasp system operates as a dynamical system at all^35,36,51^ – have remained unclear. Here, we used state space geometry analysis, dynamical systems analysis, and the LDR model with a large existing reach-to-grasp dataset to better understand the grasp control system in M1 and F5. Consistent with previous work^36^, we first found that a large common signal existed in dimensions that form a modest angle to the dimensions containing condition-specific activity. Second, we found that prior to object contact, these areas’ activity was well described by a smooth dynamical system, but after contact it was not. Third, we found that the data in M1 and F5 were better described by the LDR model than by previous models. Specifically, the key postulates of the LDR model all held in M1 and F5 during grasping. As in reaching, grasp-related activity exhibited a small set of conserved rotational frequencies in the population activity, but the planes in which the rotations existed varied systematically by condition; and the state space location and the orientation of the rotational planes were linearly related to one another and to the entire sequence of kinematics. This enabled us to use the state space location and rotation orientations to decode behavioral activity, or more interestingly, to reconstruct neural activity from kinematics. These results suggest an overall picture of the dynamics of the grasp system that is qualitatively similar to those in reaching, with the primary differences being merely quantitative: the state space location varied less for grasping than for reaching, and this was reflected in smaller angles of variation for the rotational planes for grasping than for reaching.

These quantitative differences may echo the computational demands imposed by the divergent structure and function of the arm and hand. Reaching requires controlling a small number of muscles and joints, but two reaches can require entirely different patterns of muscle activations^52,53^. Grasping, in contrast, requires coordination of several times as many muscles and joints^43,54^; but despite the requirement for high-dimensional control of the hand^43,55^, these many dimensions are smaller modulations on top of a dominant dimension of open-close^43,56–59^. Grasps for different objects therefore still require strongly correlated muscle activation patterns. These demands would be consistent with the observed differences in LDR for grasping versus what was previously shown in reaching^10^. Even more than reaching, grasping requires high-dimensional control, which is potentially implemented by the high-dimensional variation in the rotational planes. Unlike reaching, grasping may not need many completely-orthogonal planes for the dynamics because most grasps are correlated in their dominant open-close structure.

Relatedly, the similarity of the subspaces containing the common trajectory and the condition-specific activity may stem in part from the nature of the task demands. It is possible that grasp, like reach, is initiated by a condition-independent triggering signal^42^. In the present data, though, the reach locations were highly similar for all objects, and the grasps all involve an open-close cycle. These across-condition similarities might introduce the need for a strong output signal (and potentially supporting output-null activity^60^) that is the same for all the conditions tested here, even if those dimensions are capable of tuning for other reach locations or kinds of hand tasks (such as sign language^61,62^). Alternatively, this finding may reflect a fundamental difference in the system’s geometry for grasp vs. reaching. Future work could introduce more varied conditions to determine which hypothesis is correct.

The LDR model for grasp generation is compatible with many previous findings about the grasp system. Hand M1 and F5 are extensively tuned for object^63,64^, which is consistent with the state space location (which determines average tuning) depending on the object. The linear decodability of M1 for grasp^38,65–67^, including our results here, reflects a presumably large number of output-potent dimensions^60^ needed to drive the many dimensions of control. Indeed, the presence of a large number of output-potent dimensions could explain why LDR-based decoding only modestly outperformed direct decoding: LDR-based decoding benefits primarily from using output-null dimensions to denoise, and that may be less of an advantage in grasping as in reaching if the output-potent space is relatively larger in grasp. Nevertheless, the stronger description of these areas as dynamical over representational^33^ and their similarity to RNN models^39,68,69^ presumably stems from being a dynamical system, if not a conventional one. Finally, the strong deviation from being an autonomous dynamical system once object contact occurs makes sense in light of the strong sensory feedback into these areas^70–72^ that diverges from dynamical predictability with external forces^30,73,74^.

Although the finding of dynamics here agrees with some previous work^39,51^, other work has found that grasp-related activity was incompatible with dynamical systems^35^. The reason for the disagreement is not entirely clear, but there are many differences between the datasets that might explain it. A first such difference is that the present data include a reach, which the Suresh et al. work did not; there, the object was placed in the monkey’s hand by a robot and grasped to resist a pull. It is possible that in the present dataset the presence of a reach, the small variation in reach location to different objects, or the much greater involvement of the wrist add a dynamical component that was absent in the Suresh et al. dataset. Second, in the present data the movement was performed relatively unencumbered and in the dark, while Suresh et al. had the monkey’s wrist strapped down and performed the task in the light. Sensory feedback from the wrist or tendons pulling against the wrist strap, or even from continued visual input might introduce external inputs that broke the transient autonomy of the dynamical system there. Third, the present data used a much larger variety of objects and recorded more neurons, though to us neither factor seems likely to be sufficient for observing such a large qualitative difference. Fourth, the recordings here primarily sampled deeper cortical layers, leaving open the possibility that superficial layers exhibit different dynamical organization and/or stronger input dependence. Because superficial and deep layers differ in their connectivity and contributions to inter-areal communication^75,76^, future laminar recordings will be important for determining how conserved temporal structure and rotational reorientations are distributed across cortical microcircuits.

The dynamical organization observed here likely reflects the operation of a distributed sensorimotor circuit rather than motor cortex in isolation. Grasp-related activity in the motor cortex may inherit structure from upstream premotor, parietal, or somatosensory areas that encode object features, hand shape, and sensory consequences of movement^63,77–80^, with the motor cortex transforming these inputs into appropriate motor commands. Thalamocortical interactions may also play an active role in shaping cortical dynamics, consistent with emerging views of the thalamus as an active participant in motor control rather than a passive relay^81,82^. Under some circumstances it may therefore appear that a single area obeys transiently autonomous dynamics, while under other circumstances it may not.

Finally, we note several limitations of the present work. Because a reach preceded the grasp, we did not have the ability to separate grasp-related activity from reach-related activity. This is particularly relevant because grasp- and reach-related activity are thought to be mixed in single neurons^83^, consistent with general findings of mixed “selectivity”^14,84–86^ and mixed joint activations^61,67^ in single neurons throughout motor areas. Second, as discussed above, the conserved reach location across objects and the common open-close motif of grasping preclude our separating any true condition-independent trigger signal from activity that would be tuned if the movements varied in the relevant ways. Third, this model describes grasp data, but steady holding^87,88^ and force modulation^89,90^ likely require substantially different temporal structure in their control signals and are understood to be encoded less strongly in motor cortex^91^. The LDR model therefore likely would not describe these aspects of hand control. Finally, although we achieved moderately better encoding and decoding performance than with standard decoding methods, the performance of both remained far from perfect. This may stem from lower trial counts than in previous reaching data; from the shorter time period of neural activity available with this behavior; from the smaller angular range of some of the joints; or from the necessity to include data from the hold epoch (and therefore sensory feedback) in order to have enough data to analyze with encoding and decoding models. Yet another possibility is a simple numerical challenge: the higher dimensionality of movement challenges decoding models because each joint must correspond to a smaller fraction of the variance, and challenges encoding models because higher dimensionality of predictors invites overfitting. Yet, it may be that the LDR model also misses important structure – dynamical or not – in grasp activity.

In summary, our findings suggest that grasping is not an exception to dynamical systems accounts of motor control. Instead, grasp and reach appear to share a common dynamical strategy of conserved temporal structure and reorienting rotational subspaces. The principal differences are simply in the extent of location and orientation variation. These results hint that diverse forelimb motor behaviors may emerge from common dynamical building blocks whose geometric organization is adapted to the computational demands of different effectors.

## METHODS

### Subjects and behavior

#### Reach-to-grasp task

All data analyzed in this study were previously collected. Procedures for the grasping data collection are briefly described here; full details are described previously^38^. Animal care and experimental procedures followed established protocols and complied with relevant German and European regulations. Two macaque monkeys (*Macaca mulatta*; animal Z, female; animal M, male) participated.

Monkeys were trained to perform a delayed reach-to-grasp-and-hold task. After training and headpost implantation, floating microelectrode arrays were implanted in anterior intraparietal area (AIP), ventral premotor area (F5), and primary motor cortex (M1). Neural activity and hand kinematics were recorded simultaneously during task performance and analyzed offline.

The monkeys were seated in a custom chair with their heads fixed. Objects of various shapes and sizes were presented 25 cm in front of them at chest level. All objects massed 120 g regardless of size or shape. In addition to the other objects, the monkeys performed both precision and power grips on a specially designed handle.

Each trial began when the monkey pressed and held a home button and fixated on a red LED. After a variable fixation period (500–800 ms, mean: 650 ms), a spotlight illuminated the object (cue epoch, 700 ms). The spotlight turning off for a delay period (600–1000 ms, mean: 800 ms), after which the fixation LED blinked, signaling the monkey to grasp and lift the object (movement epoch) and hold it for 500 ms (hold epoch) to receive a liquid reward. Trials were immediately aborted if performed incorrectly. For the graspable handle, additional yellow or green LEDs instructed the animals to perform precision or power grips, respectively. All objects and grip types were presented in pseudorandom order, with each object successfully grasped at least 10 times before the turntable containing a set of six objects was changed. Successful, consistent-movement trial counts ranged from 7-23 per object/grasp.

#### Maze reach task

For a small number of comparison analyses, we reanalyzed previously collected neural recordings from the “maze” center-out delayed-reaching task^27^. All animal care and experimental procedures were approved by the Stanford University Institutional Animal Care and Use Committee and have been described previously. Only recordings from monkey J (*Macaca mulatta*, male) were used in the present study.

The monkey performed the task for juice reward. At the start of each trial, the monkey fixated a central point while holding a cursor floating above the fingertips at the same location on a vertical display. A target then appeared and jittered slightly during a variable instructed delay (0-1000 ms). The go cue was indicated by the cessation of target jitter, filling of the target, and disappearance of the fixation point, at which time the monkey executed a rapid reach to the target. On a subset of trials, virtual barriers were presented simultaneously with the target, requiring curved reaches around the obstacles. Together, straight and curved reaches produced 72 reaching “conditions” (36 straight and 36 curved). Neural activity was recorded simultaneously from dorsal premotor cortex (PMd) and primary motor cortex (M1) using chronically implanted 96-channel Utah arrays (Blackrock Microsystems, Salt Lake City, UT) with one implanted in each area. Both single-units and stable multi-units were included for analysis. Neural data from the maze reach task were analyzed using the preprocessing described previously^10^. The hand kinematic processing and neural preprocessing procedures detailed below were applied only to the reach-to-grasp dataset, not the maze reach dataset.

### Reach-to-grasp hand kinematics

Hand and arm kinematics were recorded using a custom glove-based electromagnetic tracking system^72^, which tracked the 3D positions of the distal interphalangeal (DIP), proximal interphalangeal (PIP), and metacarpal-phalangeal (MCP) joints for all fingers, as well as the 3D position and orientation of the hand. A wrist sensor provided forearm orientation, which was used to determine the 3D position of the elbow. The shoulder position was considered fixed due to the head restraint. This setup provided full-arm kinematics (18 joints, 27 degrees of freedom) with a temporal resolution of 100 Hz.

Kinematic joint angles were temporally binned with a 10 ms interval and smoothed using a Gaussian kernel with a standard deviation of 10 ms. Kinematic activity was aligned to the time of object contact, extending from 400 ms before contact to 300 ms after. Joint angles with limited variation were excluded based on range of motion (ROM): for each joint, the maximum ROM across condition-averaged trajectories was computed, and joints with a maximum ROM <15° were discarded. To remove atypical trials, pairwise root-mean square error (RMSE) and Pearson correlation were computed between the joint angle trajectories of all trial pairs for each condition and joint angle separately. Each trial was assigned its mean RMSE and mean correlation relative to all other trials within the same condition. Trials with RMSE z-scores >2.5 or correlation z-scores <-2.5 were classified as outliers. If fewer than five trials remained for a given joint angle within any condition, that joint angle was excluded for all conditions. Finally, any trial identified as an outlier for one or more retained joint angles was removed from all subsequent kinematic and neural analyses.

### Reach-to-grasp neural recordings

Following recovery and training with head fixation, six floating microelectrode arrays (FMAs; MicroProbes for Life Science) were implanted, with two 32-electrode arrays placed in each of areas AIP, F5, and M1. Implants were in hand-related parts of each area. Electrode lengths ranged from 1.5 to 7.1 mm^65^. Surgical procedures and perioperative care followed previously described protocols.

Spiking activity was recorded from 192 channels across AIP, F5, and M1. Data included both single-unit and multiunit activity, sampled at 24 kHz with 16-bit resolution. Spikes were detected and sorted using automated clustering followed by manual curation.

Neural activity was temporally binned with a 10 ms interval and smoothed using a Gaussian kernel with a standard deviation of 30 ms. AIP activity was aligned to the onset of the cue, from 150 ms before to 600 ms after. F5 and M1 activity were aligned to the time of object contact, extending from 400 ms before contact to 300 ms after except where noted that post-contact activity was excluded. These lockings were chosen to capture peak task-related neural modulation. Units with low firing rates were excluded based on activity within these analysis windows: for each unit, the trial-averaged firing rate was computed using the relevant event locking, and units whose maximum firing rate did not exceed 7 spikes per second were discarded. To further exclude poorly isolated or highly variable units, we computed a signal-to-noise ratio (SNR) for each neuron. For each condition, a PETH was obtained by averaging across trials. Signal amplitude was quantified as the range of firing rates across all conditions and time points in the PETH. Noise was quantified as the maximum SEM across conditions and time points. The SNR was defined as the ratio of signal amplitude to noise amplitude^42^. For M1 and F5, we excluded units with SNR values <2 from further analysis; for AIP, due to the smaller number of modulated units available, we only excluded units with SNR values <1.

### Principal Component Analysis (PCA)

For Figures 1e-f and 2i, the neural data were averaged across trials within condition, then organized in a matrix of size *CT* × *N*, where *C* denotes the number of conditions, *T* the number of time points, and *N* the number of neurons. Each neuron’s activity was soft-normalized by dividing by its firing rate range plus a small constant, chosen to be 2 spikes/s, to reduce the influence of high-firing-rate neurons while still permitting them more influence than low-rate neurons. Kinematic data were analogously reshaped into matrices of size *CT* × *J*, where *J* denotes the number of joint angles. PCA was then performed separately on each dataset. The top three principal components were retained for visualization.

### Demixed Principal Component Analysis (dPCA)

Demixed principal component analysis (dPCA) was applied to trial-averaged neural population activity to separate variance associated with time, condition and their interaction^45^. Each neuron’s activity was first soft-normalized as above. For each cortical area, neural activity was then organized into a three-dimensional tensor of size neurons x condition x time, with activity averaged across similar trials. Analyses used the standard dPCA implementation provided by the original authors.

Neural activity was decomposed into components corresponding to:

1. Condition-independent (time-only) activity
2. Condition-specific activity
3. Condition-time interaction activity

Following previous work, groups 2 and 3 were considered together as condition-specific^45^.

The primary analyses reported in this study used the standard regularized dPCA implementation without cross-validated optimization of the regularization parameter. To assess the robustness of the results, we additionally repeated the analyses using regularized dPCA with the regularization strength optimized separately for each cortical area. The cross-validated analyses produced qualitatively similar results to the primary analyses and therefore did not alter the conclusions.

To evaluate dimensionality, the number of retained components for each marginalization was varied and cumulative variance explained was computed. Because higher-order components captured comparatively little additional variance, subsequent analyses were restricted somewhat arbitrarily to the first 15 dPCA components, which together accounted for the majority of explainable variance across cortical areas. Retaining additional components did not qualitatively alter the results.

### Canonical Correlation Analysis (CCA)

To quantify the similarity of the common trajectory present in neural dynamics across cortical areas, we performed CCA on the condition-mean signal extracted from M1 compared to F5. As for other analyses, the condition-mean was computed on the PETHs for each time point independently. Because the number of recorded neurons differed across areas, and to reduce dimensionality, PCA was applied separately to the condition-mean matrix from each area. The first 10 principal components were retained, producing low-dimensional representations of the dominant common trajectory of the population dynamics. CCA was then performed on these PCA-reduced common trajectories from M1 and F5. Canonical correlation coefficients were computed between the resulting canonical variates and used to quantify the degree of shared structure in the common trajectory between areas. Higher canonical correlations indicate greater similarity in the shapes of the common trajectories expressed across cortical regions. To assess whether the observed correspondence between M1 and F5 exceeded that expected by chance, we generated a shuffle distribution (1,000 shuffles) by independently randomly permuting the temporal order of the condition-mean for each F5 neuron before PCA. PCA was then repeated on each shuffled F5 population, and the scores from the first 10 principal components were retained. CCA was subsequently performed between the unshuffled M1 representation and each shuffled F5 representation. The resulting distribution of canonical correlations provided a null against which the correlations obtained from the unshuffled data were compared. Shuffling the F5 scores after the PCA step instead produced a similar null distribution.

### jPCA

To assess rotational population dynamics, neural activity was analyzed using jPCA^5^. Analyses were performed separately for each monkey, cortical area, and grasp condition.

Following the procedures of the original work, trial-averaged neural activity was first soft-normalized as above. Activity was then mean-centered across conditions at each time point, and neural population activity was projected onto the top six principal components before applying jPCA.

Neural activity was projected onto the plane corresponding to the largest imaginary eigenvalues to visualize the dominant rotational dynamics. Model performance was quantified using the coefficient of determination (R^2^), which measures how well the fitted skew-symmetric dynamical system predicts the temporal derivatives of the neural population state.

### Fitting linear dynamical systems to single conditions

Condition-specific neural dynamics were analyzed by fitting an LDS independently to the neural population activity for each grasp condition (Fig. 3b), following previously described methods^10^. For each condition, trial-averaged neural population activity was soft-normalized as above then projected onto the first *k* principal components to obtain a reduced-dimensional neural state, *x*(*t*). Dynamics were modeled using the mapping formulation of a linear time-invariant system:

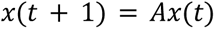

where *A* is the dynamics matrix estimated by linear least-squares regression in the reduced-dimensional state space.

To determine the optimal dimensionality *k*, trials for each condition were randomly partitioned into two groups to obtain independent estimates of trial-averaged firing rates. Low-dimensional subspaces and corresponding dynamics matrices were estimated from one partition while varying latent dimensionality. Model performance was then evaluated on the held-out partition by computing the fraction of neural population variance explained. This cross-validation procedure was repeated multiple times with independent trial partitions, and the dimensionality maximizing cross-validated variance explained was selected. Optimal dimensionalities estimated from this procedure were 5 for M1 and F5 and 3 for AIP.

To avoid unrealistically slow dynamical modes, eigenvalues corresponding to long decay time constants were capped at 10s during model fitting. Model performance was quantified by simulating the fitted dynamics forward from the neural population state at the beginning of the corresponding analysis window. For M1 and F5, analyses used activity aligned to object contact (−400 to 300 ms relative to contact crossing), whereas AIP analyses used activity aligned to cue onset (−150 to 600 ms relative to cue onset). Model performance was quantified as the fraction of neural population variance explained by the simulated trajectories for each condition.

A shared LDS was also fit by estimating a single linear dynamics matrix from the concatenated neural activity across all grasp conditions. Model performance was evaluated using the same variance-explained metric as for the condition-specific LDS.

### Factorizing neural activity using temporal basis functions (the LDR model)

To identify temporal dynamics conserved across grasp conditions, neural population activity was factorized into shared temporal basis functions and condition-specific loading matrices following the approach of previous work^10^. For each condition *c*, trial-averaged neural activity was represented as a matrix:

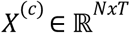

where *N* denotes the number of neurons and *T* the number of time points. Prior to factorization, firing rates were soft-normalized as above. Activity from all *C* conditions were concatenated along the neuron dimension to form a single matrix:

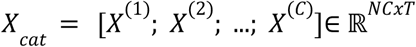

A singular value decomposition was applied to the concatenated activity matrix *X_cat_* = UΣV^T^,and the top *k* right singular vectors were taken as shared temporal basis functions: *B*∈ ℝ*^kxT^* while the corresponding left singular vectors, weighted by the singular values, were partitioned into condition-specific loading matrices *L^(c)^*∈ ℝ*^Nxk^*. For each condition *c*, neural activity was then approximated as:

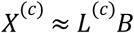

To associate temporal basis functions with interpretable dynamical modes, we fit a linear dynamical system to the temporal basis functions:

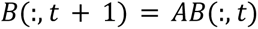

where *A* is the fitted dynamics matrix, and *B*(:, *t*) denotes the values of all temporal basis functions at time t. The temporal basis functions were then projected into the eigenvector basis of *A*. The inverse transformation was applied to each condition’s loading matrix, preserving the factorization and reconstruction accuracy while associating each temporal component with a specific dynamical mode. For most analyses, we used these ‘purified’ *L*^(*c*)^ matrices; the exception was for the encoding and decoding analyses, where using an orthogonal basis improved fitting robustness.

The dimensionality *k* of the temporal basis was determined using cross-validation. Trials were randomly partitioned into training and test sets. Temporal basis functions and condition-specific loading matrices were estimated from the training partition while varying *k*. Neural activity in the held-out partition was then reconstructed using the learned basis functions. The dimensionality maximizing cross-validated variance explained across conditions was selected.

### Subspace alignment and alignment index analysis

Subspace alignment analysis was used to assess whether condition-specific rotational dynamics occupied similar subspaces of neural state space, including the alignment index approach^48^ and related to several similar previous approaches^49,50^. Neural population activity for each condition was first projected into low-dimensional condition-specific subspaces, which were the pair of columns from the purified *L*^(*c*)^’s corresponding to the rotation plane of interest.

For a given component (i.e., the 1D offset or a 2D subspace containing a rotation), let *X_a_*∈ℝ*^Nxk^* denote the trial-averaged neural activity for condition *a*, and let *Q_b_*∈ℝ*^Nxk^* denote the matrix whose columns span the subspace associated with the same component in condition *b*. Alignment of condition *a* to condition *b* was computed as the fraction of variance in *X_a_* explained by projection onto the subspace of condition *b*:

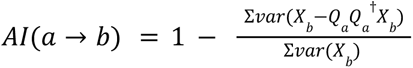

where † indicates pseudoinversion (Moore-Penrose inverse), var(·) denotes the variance across time for each neuron, and Σ*var* indicates the sum of these variances across all neurons. Alignment values range from 0 (orthogonal subspaces) to 1 (identical subspaces). This metric is asymmetric, so alignment was computed for all ordered pairs of conditions within each cortical area and each component (1D offset or 2D rotational subspace). Self-alignments (*a* = *b*) were excluded from statistical comparisons. Subspace alignment (e.g., Fig. 2f,g) quantifies, for each dimension, the fraction of variance captured by the corresponding dimension, complementing the alignment index analysis.

To estimate the contribution of finite sampling variability, we constructed a control distribution using split-half resampling of single-trial data within each condition. For each resampling iteration, trials were randomly partitioned into two halves, trial-averaged activity was computed separately for each partition, and independent subspaces were estimated from each half using the same projection procedure as in the main analysis. Alignment indices were then computed between subspaces derived from the two estimates of the same condition. This procedure was repeated 100 times to obtain a distribution of within-condition alignment values, which served as a noise baseline for comparison against across-condition alignment.

### Subspace Excursion Angles (SEAs)

We quantified the geometry of condition-specific rotational neural subspaces in high-dimensional neural population space using Subspace Excursion Angles (SEAs), following the conceptual framework and algorithm previously described^10^. For each rotational component, each condition was associated with a two-dimensional loading subspace 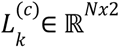, the two columns of the purified *L*^(*c*)^ corresponding to the embedding of the two-dimensional rotational plane in neural state space.

SEAs were computed iteratively using principal angles between subspaces. First, all pairwise principal angles between subspaces were computed. The pair of subspaces exhibiting the largest principal angle defined the initial step of the sequence. One of the two subspaces and the vector in the second subspace forming that largest principal angle became the reference. The process of finding the subspace producing the largest principal angle relative to the reference was repeated, and the sequence was extended accordingly. The first SEA therefore quantified the largest angle between any pair of subspaces, while subsequent SEAs quantified additional maximal angles not included in the accumulated reference subspace. For each rotational component and cortical area, the first 40 SEAs were computed. The largest possible sequence of SEAs was found. All procedures were identical for the state space location, except that the subspaces were in ℝ^*Nx*1^.

To assess the contribution of estimation noise, a control distribution was constructed using the same split-half trial procedure described above. For each condition, independent subspaces were estimated from two non-overlapping trial partitions, and SEAs were computed on these paired estimates. Observed SEAs across conditions were then compared against this within-condition control distribution.

### Predicting rotation orientations from other rotations or the state space location

To test whether the parameters of one rotational mode predicted those of another, we used a previously-described^10^ cross-validated decoding approach based on the temporal basis factorization. For each cross-validation split, temporal basis functions and condition-specific loading matrices were estimated from the training conditions. Each rotational mode was represented by the pair of (purified) loading matrix columns corresponding to its two-dimensional rotational plane. For each pair of rotational modes, the predictor loading matrix columns were vectorized across neurons, dimensionality was reduced with SVD to retain 80% of the variance, and ridge regression (λ=0.1) was used to predict the vectorized loading matrix columns of the target rotational mode across conditions. Results were robust to the choice of regularization parameter over λ∈[0,1]. Model performance was evaluated using 4-fold cross-validation across conditions. For each held-out condition, the predicted loading matrix columns were reshaped to their original dimensions and multiplied by the corresponding temporal basis functions to reconstruct the predicted rotational component of the neural activity. Prediction accuracy was quantified as the variance explained between the predicted and observed rotational component for each held-out condition. Statistical significance was assessed using a permutation test with 10,000 iterations. For each permutation, rotational components were independently reassigned across conditions prior to reconstruction of the neural activity, preserving the temporal basis functions while disrupting the correspondence between rotational modes. The complete decoding analysis was repeated on each shuffled dataset to generate a null distribution of mean variance explained for every predictor-target pair. Empirical one-sided permutation p-values were computed as the proportion of shuffled datasets whose mean variance explained equaled or exceeded that obtained from the unshuffled data.

To determine whether the state-space location predicted the rotational structure, the loading matrix column corresponding to the state-space location was used as a predictor, dimensionality was reduced as above, and the rotation loadings were predicted using ridge regression (λ=0.01). Results were robust to the choice of regularization parameter over λ∈[0,1]. Model performance was evaluated using leave-two-out cross-validation across conditions. Prediction accuracy and statistical significance were quantified as above.

### Analysis of one vs. both dimensions of rotational planes changing

We quantified whether both dimensions of rotational planes vary or whether only one changes while the other is preserved. For each rotation, we first orthonormalized 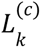. Principal angles between pairs of condition-specific subspaces (the 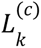 for different *c*’s) were then computed. Because each rotational plane is two-dimensional, this procedure yields two principal angles per condition pair, corresponding to the smallest and largest angles between subspaces, θ_1_ and θ_2_.

### Tangling

To quantify the degree to which neural population dynamics followed smooth, autonomous trajectories, we computed the tangling metric as previously described^28^. Tangling quantifies how strongly differences in neural state relate to differences in instantaneous state derivatives, with higher values indicating less consistent (more input-driven) dynamics.

For each cortical area, trial-averaged firing rates were soft-normalized across neurons, mean-centered, and concatenated across conditions. Neural activity was then projected into a low-dimensional state space using PCA. The first 10 principal components were retained, capturing the majority of population variance; results were robust to alternative dimensionalities.

Tangling between two time points *t* and *t*’ was defined as:

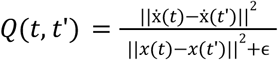

where ɛ is a small constant added for numerical stability. For each time point, the tangling index was defined as:

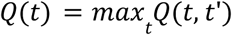

Analyses were performed across all conditions and using identical preprocessing and dimensionality reduction procedures for neural and kinematic data to enable direct comparison, except that joint-angle trajectories were range normalized instead of soft-normalized.

### Encoding

Encoding performance was evaluated using five-fold cross-validation over conditions. Conditions rather than individual trials were assigned to folds, such that all trials from a given condition were withheld together during testing. Within each training fold, neural and kinematic variables were independently normalized using training-set statistics. The resulting means and standard deviations were subsequently applied to the held-out test data.

#### Linear (direct encoding)

Linear encoding models were used to predict neural population activity from kinematic variables. For each cross-validation fold, ordinary least-squares regression was fit using normalized kinematics as predictors and normalized neural activity as responses:

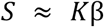

where *S* denotes (spiking) neural activity, *K* denotes kinematic variables, and β is the encoding weight matrix fit using least-squares regression. The resulting model was applied to held-out kinematic data to generate predicted neural activity. Similar results were obtained with ridge regression.

#### Location-Dependent Rotations (LDR)

We implemented LDR encoding following the method introduced previously^10^. Within each training fold, soft-normalized neural activity was reshaped into a matrix of size neurons * trials by time and factorized using truncated SVD, and grouping the singular values into the left singular vectors to make a loadings term:

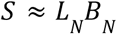

where *S* is the (spiking) neural activity, *L*_N_ contains the neural loading coefficients and *B_N_* contains the neural temporal basis functions. The decomposition was truncated to *k*_N_ latent dimensions.

Kinematic activity was decomposed analogously:

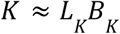

where *L*_K_ are the kinematic feature coefficients and *B*_K_ contains the kinematic basis functions. The decomposition was truncated to *k*_K_ dimensions.

Loadings were reshaped such that each trial was represented by a single latent-state vector. A mapping between kinematic and neural latent spaces was then estimated from training data:

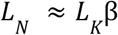

where β is the latent-space transformation matrix fit using least-squares regression.

For held-out conditions, normalized kinematic activity *K*^(*test*)^ was projected onto the training-derived kinematic basis traces to obtain latent kinematic coefficients:

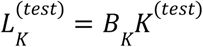

Predicted neural latent coefficients were then computed as:

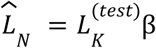

Predicted neural activity was reconstructed by projecting the estimated neural latent coefficients through the neural temporal basis functions:

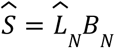

#### Selection of latent dimensionality

Neural latent dimensionality (*k*_N_) and kinematic latent dimensionality (*k*_K_) were systematically varied from 1 to 19 in increments of 2, and for each dimensionality pair five-fold cross-validated encoding performance was computed independently for each neuron. The dimensionality pair yielding the highest mean cross-validated performance was identified for each neuron, and the final dimensionalities used for population analyses were selected as the optimal neural and kinematic dimensionalities across neurons.

#### Performance metric

Encoding performance was quantified using the coefficient of determination computed on condition-averaged peri-event time histograms (PETHs). For each cross-validation fold, predicted neural activity for held-out trials were averaged across repetitions of each test condition to generate predicted PETHs. Performance was calculated independently for each neuron as:

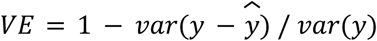

where *y* and *ŷ* denotes observed and predicted PETHs values, respectively. Reported encoding performance corresponds to the mean held-out PETH variance explained across cross-validation folds.

### Decoding

Decoding performance was evaluated using five-fold cross-validation. Within each training fold, neural and kinematic variables were independently normalized using training-set statistics. The resulting means and standard deviations were applied to the held-out test data. Predicted kinematics were transformed back into physical units prior to performance evaluation.

#### Linear direct

Linear decoding was performed using ordinary least-squares regression to predict kinematics from neural population activity. Predictions were converted back to physical units using the training-set normalization parameters.

#### Nonlinear direct

To evaluate nonlinear relationships between neural activity and behavior, we used support vector regression with a nonlinear kernel. An independent decoder was trained for each joint variable using the neural population activity as input. Models were implemented using MATLAB’s fitrsvm function with Gaussian kernels, automatic kernel-scale selection, and a box constraint of 1. Because neural activity was normalized before model fitting, additional internal standardization was disabled. For each fold, trained models were applied to held-out neural activity to generate decoded kinematic trajectories, which were subsequently transformed back into physical units.

#### Linear LDR

We implemented LDR decoding following the framework introduced previously^10^. Encoding models were fit as described above, then the relationship was pseudoinverted to permit decoding the kinematics loadings from the neural loadings:

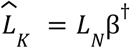

where † denotes the pseudoinverse.

Decoded kinematic trajectories were reconstructed by projecting the predicted latent coefficients through the kinematic basis traces:

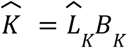

Held-out neural activity was projected onto the neural temporal basis functions derived from the training data before latent decoding.

#### Nonlinear LDR

To determine whether nonlinear transformations between neural and kinematic latent spaces improved decoding performance, we replaced the linear latent mapping with support vector regression with a nonlinear kernel, as for nonlinear direct decoding. All other steps were identical to linear LDR decoding.

#### Performance metric

For both LDR approaches, neural latent dimensionality (*k_N_*) and kinematic latent dimensionality (*k_K_*) were optimized as for the encoding models. Decoding performance was quantified as the fraction of variance explained (VE) computed in the original kinematic coordinate system:

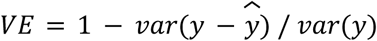

where *y* and *ŷ* denotes observed and predicted kinematic values, respectively. VE was computed independently for each joint angle and cross-validation fold. Reported decoding performance corresponds to the mean held-out VE across folds.

### Comparison of state-space location distances and rotational diameters

For the analyses in Figure 7, to quantify location differences between conditions we analyzed the first temporal mode. For each pair of conditions (*i*, *j*), a difference trajectory was computed as:

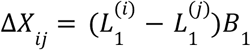

The Euclidean norm of the difference trajectory was evaluated at each time point, and the 90^th^ percentile across time was taken as the location distance for that condition pair.

To quantify the state space diameter of the rotational structure, we analyzed the subspace defined by the first oscillatory mode. For each condition, the reconstructed oscillatory trajectory was computed as:

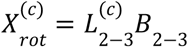

The reconstructed rotational trajectory was projected onto its first two principal component axes to define a 2-D orthogonal coordinate system spanning the rotational plane. Let *F*^(*c*)^∈ℝ^2×*T*^ denote the resulting trajectory. Rotational diameter was estimated by projecting the trajectory onto unit vectors spanning angles from 0° to 360°. For each projection direction, the range of projected values across time was computed, and the maximal range across all directions was taken as the rotational diameter:

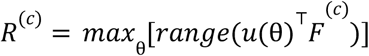

where *u*(θ) is a unit vector oriented at angle θ.

For visualization, distributions of pairwise location distances and condition-specific rotational diameters were displayed using kernel density estimates. To compare rotational and translational scales, a diameter-to-location ratio was computed from the data. Uncertainty in this ratio was estimated using bootstrap resampling (5,000 iterations). At each iteration, rotational diameters and location distances were independently resampled with replacement, and the median of the resulting diameter-to-location ratio was computed. The resulting bootstrap distribution was used to estimate 95% confidence intervals and p-values for the ratio.

## Notes

### Competing Interest Statement

The authors have declared no competing interest.

